# Exploring the therapeutic potential of defective interfering particles in reducing the replication of SARS-CoV-2

**DOI:** 10.1101/2024.03.11.584367

**Authors:** Macauley Locke, Dmitry Grebennikov, Igor Sazonov, Martín López-Garcá, Marina Loguinova, Andreas Meyerhans, Gennady Bocharov, Carmen Molina-París

## Abstract

SARS-CoV-2 still presents a global threat to human health due to the continued emergence of new strains and waning immunity amongst vaccinated populations. Therefore, it is still relevant to investigate potential therapeutics, such as therapeutic interfering particles (TIPs). Mathematical and computational modelling are valuable tools to study viral infection dynamics for predictive analysis. Here, we expand on the previous work by Grebennikov *et al*. (2021) on SARS-CoV-2 intra-cellular replication dynamics to include defective interfering particles (DIPs) as potential therapeutic agents. We formulate a deterministic model that describes the replication of wild-type (WT) SARS-CoV-2 virus in the presence of DIPs. Sensitivity analysis of parameters to several model outputs is employed to inform us on those parameters to be carefully calibrated from experimental data. We then study the effects of co-infection on WT replication and how DIP dose perturbs the release of WT viral particles. Furthermore, we provide a stochastic formulation of the model that is compared to the deterministic one. These models could be further developed into population-level models or used to guide the development and dose of TIPs.

**Author summary:** SARS-CoV-2 continues to evolve, with new strains or sub-strains being identified thanks to efforts to monitor the virus. Consequently, new strains threaten human health as current vaccinations may not adequately protect against future strains. It is therefore important to understand the roles that additional therapeutics could play in protecting against these future strains.

Therapeutic interfering particles (TIPs), otherwise referred to as defective interfering particles (DIPs), could provide an additional treatment option against future strains. Previous models have examined the role of DIPs at the within-host level during co-infection with wild-type virus, but have paid little attention to intra-cellular dynamics. Here we extend the previous intra-cellular replication model of SARS-CoV-2 by Grebennikov *et al*. (2021) to include co-infection of WT virus with DIPs. We show that DIPs lead to a reduction in the WT virus in a dose-dependent manner, with higher doses leading to up to 10-fold reduction in total WT virus released from a cell depending on the multiplicity of infection (MOI). We find these results to be consistent for both deterministic and stochastic formulations of the model. Our approaches could be developed into a within-host model or population-level model, which could then be used to guide therapeutic DIP doses.

## Introduction

In December 2019, a new infectious disease was reported to the World Health Organisation (WHO) that would later be identified as a novel coronavirus (SARS-CoV-2) [1]. By 30 January 2020, the WHO declared SARS-CoV-2 a “public health emergency of international concern” [2], as it rapidly spread to 113 countries. By the 11th of March 2020, it had caused 118,319 infections and 4,292 deaths. Consequently, the WHO declared SARS-CoV-2 a pandemic [3, 4], and as of the 29th of July 2022, about 572 million infections and over 6 million deaths have been recorded worldwide. During the early stages of the pandemic, treatment options were limited to chloroquine and remdesivir [5, 6]. However, since then several effective vaccines have been developed that provide protection and reduce transmission, with many countries rolling out mass vaccination programs [7]. Although vaccines for SARS-CoV-2 now exist, the emergence of new strains due to mutations has led to further concerns about vaccine effectiveness [8, 9]. This fact, together with that of waning immunity and the existence of individuals who are unable to be vaccinated or out-right refuse to do so, highlight the need for additional therapeutics and prophylactics [10, 11].

One such potential therapy is viral interfering particles. During viral replication, mutants lacking essential parts of the viral genome arise [12, 13], which are unable to replicate in the absence of wild-type (WT) virus. These are known as defective interfering particles (DIPs). DIPs can be exploited to make therapeutic interfering particles (TIPs), which inhibit the replication of WT virus by outcompeting WT gene segments for resources required during viral replication and assembly [14, 15]. TIPs/DIPs have been investigated for several viruses, including HIV, Ebola, influenza, and SARS-CoV-2 and have been found to cause a two-fold reduction in viral titres [14–16]. However, caveats exist in their production; for instance, which sections of the viral genome are to be removed to allow for replication at a faster rate than WT, they are virus-specific, and little is known about how mutations change replication dynamics [13, 17].

From a mathematical modelling perspective, a long-standing effort exists to describe transmission dynamics at the population and within-host levels (see Ref. [18] and references therein). At the within-host level DIPs, as therapeutics, have been studied in Refs. [19, 20]. However, little effort has been devoted to investigating the intra-cellular replication kinetics of DIPs in the presence of WT virus. Grebennikov *et al*. [21] have recently proposed a SARS-CoV-2 intra-cellular replication dynamics model. This model allowed for the quantification of viral genomes and proteins during the replication cycle. We wish to exploit this model to explore co-infection with DIPs and the effect of DIPs on the replication dynamics of the WT virus. In particular, in this study, we formulate a mathematical model of SARS-CoV-2 replication in a cell co-infected with DIPs. As in Ref. [21], we will follow a deterministic approach to calibrate model parameters. We shall use sensitivity analysis to study the impact parameters have on the release of both WT and DIP viral particles. We also introduce a stochastic description of this model to compare to the deterministic one. We shall also investigate how initial doses of each virus affect viral particle production (WT and DIPs) to quantify DIP-related inhibition of WT replication and the reliance of DIPs on the WT replication machinery.

## Materials and methods

The kinetics of the corresponding biochemical reactions are described in the deterministic mathematical model introduced in the **mathematical model** section of this paper. The system of ordinary differential equations (ODEs) is formulated under the assumption of mass action kinetics, Michaelis-Menten approximations, and on the biological scheme presented in Fig 1. The model can, in principle, be defined as a stochastic process.

**Fig 1.**
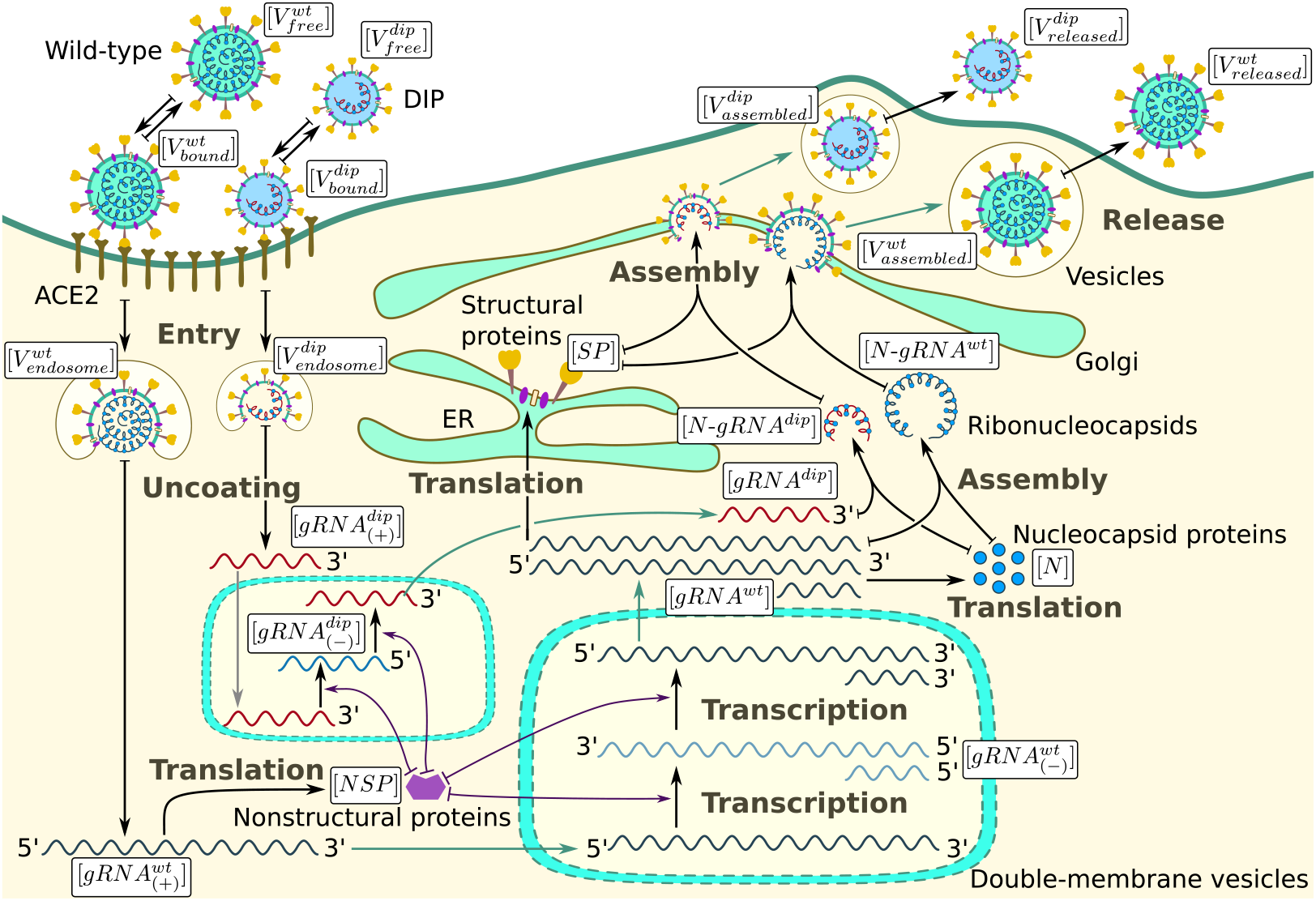
Biological scheme of the competitive replication of the infectious SARS-CoV-2 and defective interfering viral particles. There are three types of arrows shown on the scheme: (i) arrows without a T-end (which are not green) indicate the synthesis processes, i.e., translation or transcription, (ii) arrows with T-shaped beginning indicate that the variable at which the arrow points with the arrow-end is increased while the variable at which the arrow points with a T-end is decreased (e.g., transitions of the entities from one state to another, or a decrease of non-structural proteins during the transcription activation), (iii) green arrows indicate the transport of the entities from one place to another (to or from double-membrane vesicles or to the cell membrane in vesicles), which is not modelled explicitly. All entities are subject to degradation, however, these processes are not shown to avoid cluttering the figure.

### Sensitivity analysis

The mathematical model included parameters which encode the biological mechanisms under investigation. Since many parameters required calibration, it is important to identify which have the greatest effect on model outputs. Global sensitivity analysis allowed us to evaluate the results of simultaneous changes in parameter values [22]. For implementing this approach, consider the vector of parameters ***θ*** = (*θ*_1_, *θ*_2_, …, *θ*_*n*_) such that the model output is described as *Y* = *g*(***θ***). We use the Sobol approach to determine global sensitivities [23]. Each parameter *θ*_*i*_ can be considered a random variable with an associated range. Since *Y* is a function of these variables, it is also a random variable with variance *V* (*Y*). We were interested in the conditional variance 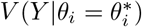. However, since the value of 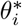 is not known, we instead considered the average conditional variance, 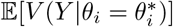, where the expectation is with respect to *θ*_*i*_, and the variance is taken over all remaining parameters *θ*_*j*_, *j*≠*i*. The law of total probability gives

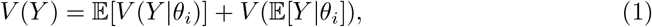

from which the first order Sobol index for parameter *θ*_*i*_ is defined as

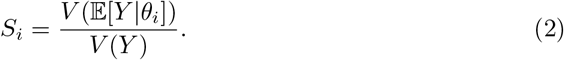

We also investigated the result of multiple fixed parameter values. If we let 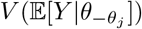 be the expected reduction in the variance by fixing all parameters except *θ*_*j*_, then the total effect of parameter *θ*_*i*_ can be defined as

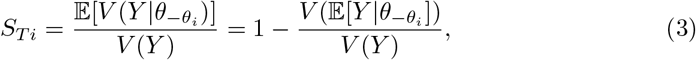

where a larger sensitivity index indicates greater importance of the associated parameter to the given model output [22].

### Model development and calibration

In formulating the new mathematical model, we introduced several additional parameters that relate to the kinetics of DIPs and the loss of non-structural proteins due to DIPs using trans-elements from WT virus for their replication. These variables are summarised in Table 1. Grebennikov *et al*. provided parameter estimates for the WT virus [21]. These values are summarised in Table 2.

**Table 1.** Dynamical variables of the mathematical model for the life cycle of SARS-CoV-2, with defective interfering particles.

| Variable | Definition | Value |
| --- | --- | --- |
| $[V_{free}^{wt}]$ | number of free infectious (*i.e.*, wild-type) virions outside the cell membrane | 10 |
| $[V_{bound}^{wt}]$ | number of infectious virions bound to ACE2<br>and activated by TMPRSS2 | 1-10 |
| $[V_{endosome}^{wt}]$ | number of infectious virions in endosomes | 1-10 |
| $[V_{free}^{dip}]$ | number of free non-infectious (*i.e.*, defective interfering particles)<br>virions outside the cell membrane | |
| $[V_{bound}^{dip}]$ | number of non-infectious virions bound to ACE2<br>and activated by TMPRSS2 | |
| $[V_{endosome}^{dip}]$ | number of non-infectious virions in endosomes | |
| $[gRNA_{(+)}^{wt}]$ | single strand positive sense genomic RNA | 1-5 |
| $[gRNA_{(+)}^{dip}]$ | single strand positive sense DIP genomic RNA | |
| $[NSP]$ | population of non-structural proteins | — |
| $[gRNA_{(-)}^{wt}]$ | negative sense genomic and subgenomic RNAs of infectious virus | 10 |
| $[gRNA^{wt}]$ | positive sense genomic and subgenomic RNAs of infectious virus | $10^4$ |
| $[gRNA_{(-)}^{dip}]$ | negative sense subgenomic RNAs of DIPs | |
| $[gRNA^{dip}]$ | positive sense subgenomic RNAs of DIPs | |
| $[SP]$ | total number of structural proteins<br>$S + M + E$ per virion | $2 \times 10^3 \in (1, 125, 2, 230)$ |
| $[N]$ | $N$ proteins per virion | $456 ; 1, 465 \in (730, 2, 200)$ |
| $[N-gRNA^{wt}]$ | ribonucleocapsid molecules for infectious virions | |
| $[N-gRNA^{dip}]$ | ribonucleocapsid molecules for non-infectious virions | |
| $[V_{assembled}^{wt}]$ | assembled infectious virions in endosomes | — |
| $[V_{released}^{wt}]$ | released infectious viruses | $10 - 10^4$ virions<br>in 7 to 24 hours |
| $[V_{assembled}^{dip}]$ | assembled non-infectious virions in endosomes | |
| $[V_{released}^{dip}]$ | released non-infectious virions | |

**Table 2.** Estimates of previously calibrated model parameter values.

| Parameter | Description, Units | Value | Range [References] |
| --- | --- | --- | --- |
| $k_{bind}$ | rate of virion binding to ACE2 receptor, $h^{-1}$ | 12 | (3.6, 12) [24, 25] |
| $d_V^{wt}$ | clearance rate of WT extracellular virions, $h^{-1}$ | 0.12 | (0.06, 3.5) [26–28] |
| $k_{diss}$ | dissociation rate constant of bound virions, $h^{-1}$ | 0.61 | (0.32, 1.08) [24, 25] |
| $k_{fuse}$ | fusion rate constant, $h^{-1}$ | 0.5 | (0.33, 1) [29] |
| $k_{uncoat}$ | uncoating rate constant, $h^{-1}$ | 0.5 | (0.33, 1) [29] |
| $d_{endosome}^{wt}$ | degradation rate of WT virions in endosomes, $h^{-1}$ | 0.06 | (0.0001, 0.12) [26, 30] , |
| $k_{transl}$ | translation rate, nt/mRNA $h^{-1}$ | 45,360 | (40,000, 50,000) [31, 32] |
| $1/f_{ORF1}$ | length of ORF1 of the RNA genome coding $NSPs$, nt | 21,000 | fixed [33] |
| $d_{NSP}$ | degradation rate of proteins in the cell, $h^{-1}$ | 0.069 | (0.023, 0.69) [32, 34],<br>tuned to (0.023, 0.1) |
| $k_{tr(-)}^{wt}$ | transcription rate of WT negative sense<br>genomic and subgenomic RNAs, copies/mRNA $h^{-1}$ | 3 | (1, 100) [32],<br>tuned to (1, 20) |
| $K_{NSP}$ | threshold number of $NSPs$<br>enhancing vRNA transcription, molecules | 100 | (10, 150) |
| $d_{gRNA}^{wt}$ | degradation rate of WT positive sense RNAs in cell, $h^{-1}$ | 0.2 | (0.069, 0.69) [32, 35],<br>tuned to (0.069, 0.4) |
| $d_{gRNA(-)}^{wt}$ | degradation rate of WT negative sense RNAs<br>in double-membrane vesicles, $h^{-1}$ | 0.1 | (0.05, 0.2) |
| $k_{tr(+)}^{wt}$ | replication rate of positive sense WT RNAs, copies/mRNA/h | 1000 | (620, 1380) [36] |
| $k_{complex}^{wt}$ | rate of the WT nucleocapsid formation [$N$-gRNA], $h^{-1}$ | 0.4 | (0.02, 0.4) [37–41] |
| $K_N$ | threshold number of $N$ proteins at which<br>nucleocapsid formation slows down, molecules | $5 \times 10^6$ | $(3.5, 6.5) \times 10^6$ [42–44] |
| $1/f_N$ | length of RNA genome coding $N$ protein, nt | 1200 | fixed [45] |
| $1/f_{SP}$ | length of genome coding structural proteins $S$, $E$, $M$, nt | $10^4$ | fixed [45] |
| $d_N$ | degradation rate of $N$ protein, $h^{-1}$ | 0.023 | (0.023, 0.069) [34] |
| $d_{SP}$ | mean degradation rate of the pool of $E$, $S$, $M$ proteins, $h^{-1}$ | 0.044 | (0.023, 0.36) [34] |
| $n_{SP}^{wt}$ | total number of structural proteins $S$, $M$, $E$ per WT virion, molecules | $2 \times 10^3$ | (1125, 2230) [37, 46, 47] |
| $n_N^{wt}$ | number of $N$ protein per WT virion, molecules | 456 | fixed [37] |
| $K_{Vrel}^{wt}$ | threshold number of WT virions at which<br>the virion assembly process slows down, virions | $10^3$ | (10, $10^4$) [42, 48] |
| $k_{asemb}^{wt}$ | rate of WT virion assembling, $h^{-1}$ | 1 | (0.01, 10) [30, 49] |
| $d_{N-gRNA}^{wt}$ | degradation rate of WT ribonucleoprotein, $h^{-1}$ | 0.2 | (0.069, 0.69) [32, 35] |
| $k_{release}^{wt}$ | rate of WT virion release via exocytosis, $h^{-1}$ | 8 | (8, 7,200) [49, 50] |
| $d_{assembled}^{wt}$ | assembled WT virion degradation rate, $h^{-1}$ | 0.06 | ($10^{-4}$, 0.12) [26] |

The remaining parameters were estimated using approximate Bayesian computation (ABC) [51]. The ABC algorithm allows a user to define a set of prior beliefs about parameter distributions, *π*(***θ***), and combine this with model simulations and data to arrive at a posterior distribution *π*(***θ***|***D***). Given a sample parameter set, ***θ***^*∗*^ *∼ π*(***θ***), a user can simulate data ***D***^*∗*^ *∼ π*(***D***|***θ***^*∗*^) and compare them to the experimental data, ***D***. If the simulated data are within a given threshold distance, *ε* (with distance measure *d*(*·, ·*)), from the experimental data, *D*, then the sample parameter set (***θ***^*∗*^, ***D***^*∗*^) is accepted. Otherwise, the parameter set is rejected and this process is continued until *N* parameter sets are accepted [51]. We made use of an Euclidean distance measure, defined as

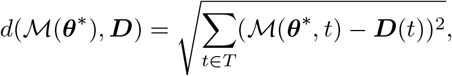

where *T* is the set of time points within the experimental data set, ***D***, and *ℳ* is the mathematical model under consideration.

Data sets for defective interfering particles are limited, with little investigation of the intra-cellular replication kinetics of WT virus in the presence of DIPs. Chaturvedi *et al*. [15] investigated two SARS-CoV-2 DIPs as TIPs. Both DIPs had shorter genomes, around 6%-10% than the WT virus. Chaturvedi *et al*. [15] performed a virus yield-reduction assay by transfecting Vero cells with TIP or control RNAs (one μg/million cells) 24 hours before infection with SARS-CoV-2 at an MOI=0.05, and harvesting culture supernatants for titration at various time-points (24, 48, or 72 hours post-infection). They discovered that these particles lead to a 1.5 *−* 1.2 log fold reduction in virus produced compared to control samples. We compared the fold reduction generated by therapeutic interfering particle two (TIP2) [15], at 24 and 48 hours, to the fold reduction from our mathematical model of 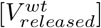 against the original model proposed by Grebennikov *et al*. [21]. These fold reductions are summarised in Table 3. For the ABC rejection method, given that the choice of a suitable *ε* is difficult, we sampled 10^6^ parameter sets and kept the top 0.1% that minimise the distance measure *d*(*·, ·*). We assumed uniform prior distributions for the parameters within the search ranges summarised in Table 4.

**Table 3.** Fold log reductions for 24 and 48 hours post-infection as reported in Ref. [15] for TIP2 to 2 decimal places (2.d.p.).

| Time (hours) | Fold log reduction (2 d.p.) |
| --- | --- |
| 24 | 1.20 |
| 48 | 1.14 |

**Table 4.**
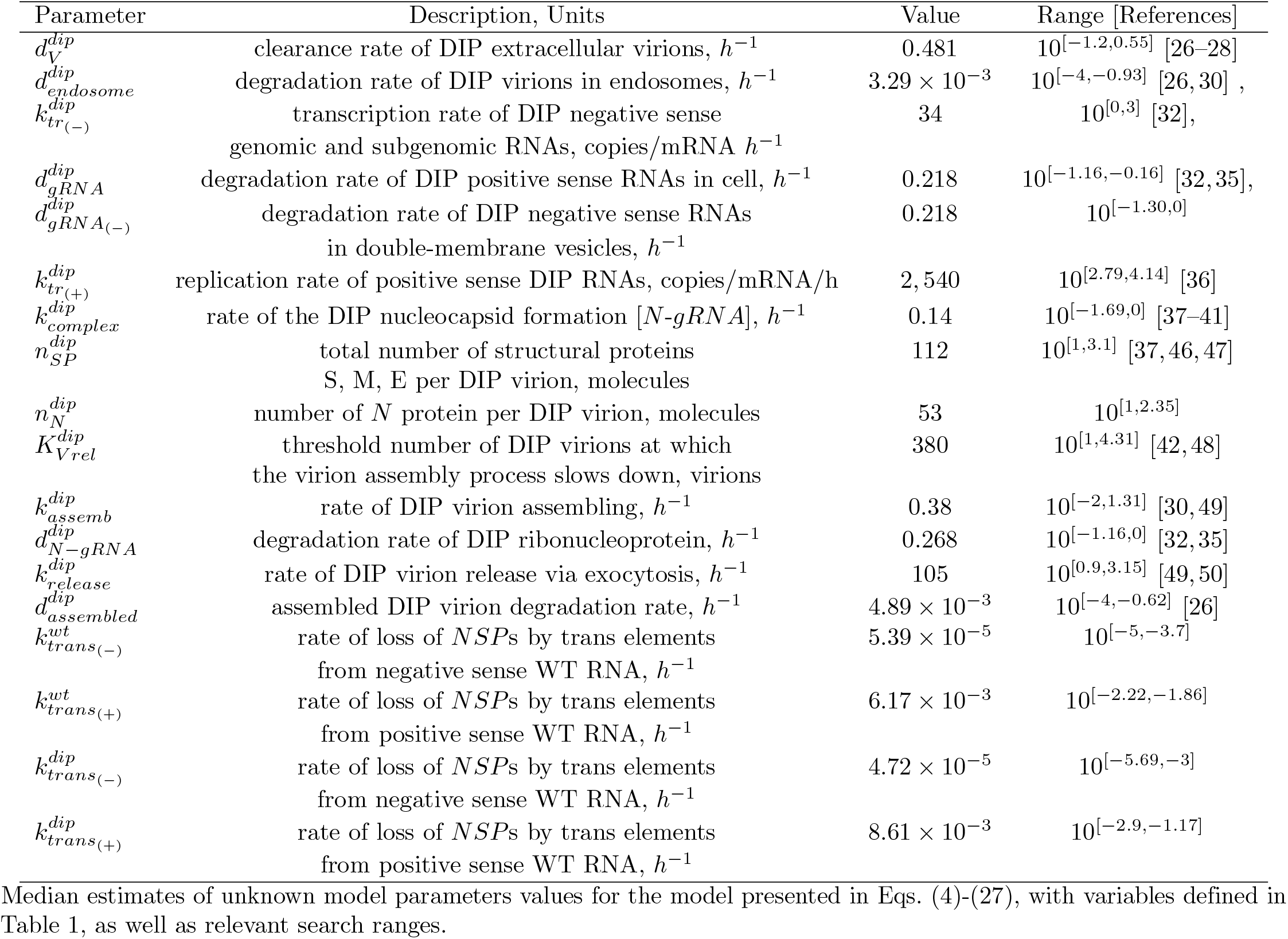
Median estimates of unknown model parameter values.

| Parameter | Description, Units | Value | Range [References] |
| --- | --- | --- | --- |
| $d_V^{dip}$ | clearance rate of DIP extracellular virions, $h^{-1}$ | 0.481 | $10^{[-1.2, 0.55]}$ [26–28] |
| $d_{endosome}^{dip}$ | degradation rate of DIP virions in endosomes, $h^{-1}$ | $3.29 \times 10^{-3}$ | $10^{[-4, -0.93]}$ [26, 30], |
| $k_{tr(-)}^{dip}$ | transcription rate of DIP negative sense<br>genomic and subgenomic RNAs, copies/mRNA $h^{-1}$ | 34 | $10^{[0, 3]}$ [32], |
| $d_{gRNA}^{dip}$ | degradation rate of DIP positive sense RNAs in cell, $h^{-1}$ | 0.218 | $10^{[-1.16, -0.16]}$ [32, 35], |
| $d_{gRNA(-)}^{dip}$ | degradation rate of DIP negative sense RNAs<br>in double-membrane vesicles, $h^{-1}$ | 0.218 | $10^{[-1.30, 0]}$ |
| $k_{tr(+)}^{dip}$ | replication rate of positive sense DIP RNAs, copies/mRNA/h | 2, 540 | $10^{[2.79, 4.14]}$ [36] |
| $k_{complex}^{dip}$ | rate of the DIP nucleocapsid formation [$N$-gRNA], $h^{-1}$ | 0.14 | $10^{[-1.69, 0]}$ [37–41] |
| $n_{SP}^{dip}$ | total number of structural proteins<br>S, M, E per DIP virion, molecules | 112 | $10^{[1, 3.1]}$ [37, 46, 47] |
| $n_N^{dip}$ | number of $N$ protein per DIP virion, molecules | 53 | $10^{[1, 2.35]}$ |
| $K_{Vrel}^{dip}$ | threshold number of DIP virions at which<br>the virion assembly process slows down, virions | 380 | $10^{[1, 4.31]}$ [42, 48] |
| $k_{assemb}^{dip}$ | rate of DIP virion assembling, $h^{-1}$ | 0.38 | $10^{[-2, 1.31]}$ [30, 49] |
| $d_{N-gRNA}^{dip}$ | degradation rate of DIP ribonucleoprotein, $h^{-1}$ | 0.268 | $10^{[-1.16, 0]}$ [32, 35] |
| $k_{release}^{dip}$ | rate of DIP virion release via exocytosis, $h^{-1}$ | 105 | $10^{[0.9, 3.15]}$ [49, 50] |
| $d_{assembled}^{dip}$ | assembled DIP virion degradation rate, $h^{-1}$ | $4.89 \times 10^{-3}$ | $10^{[-4, -0.62]}$ [26] |
| $k_{trans(-)}^{wt}$ | rate of loss of $NSPs$ by trans elements<br>from negative sense WT RNA, $h^{-1}$ | $5.39 \times 10^{-5}$ | $10^{[-5, -3.7]}$ |
| $k_{trans(+)}^{wt}$ | rate of loss of $NSPs$ by trans elements<br>from positive sense WT RNA, $h^{-1}$ | $6.17 \times 10^{-3}$ | $10^{[-2.22, -1.86]}$ |
| $k_{trans(-)}^{dip}$ | rate of loss of $NSPs$ by trans elements<br>from negative sense WT RNA, $h^{-1}$ | $4.72 \times 10^{-5}$ | $10^{[-5.69, -3]}$ |
| $k_{trans(+)}^{dip}$ | rate of loss of $NSPs$ by trans elements<br>from positive sense WT RNA, $h^{-1}$ | $8.61 \times 10^{-3}$ | $10^{[-2.9, -1.17]}$ |

### Stochastic simulation algorithm

To perform stochastic simulations of the model formulated as a Markov chain, the most popular exact approach is Gillespie’s direct method [52]. The Markov chain specifies the propensity *a*_*m*_ for the *m*-th jump process (i.e., the respective elementary reaction rate), which changes the variables by a discrete amount when that process takes place. The propensity *a*_*m*_ defines the probability *p*_*m*_ = *a*_*m*_*dt* that the *m*-th process is triggered in the infinitesimal time interval [*t, t* + *dt*). At each step of the simulation, two random numbers *r*_1_, *r*_2_ *∼ U* (0, 1) are generated to sample the time of the next jump process, *τ*, and the index *r*_*m*_ of the process to perform:

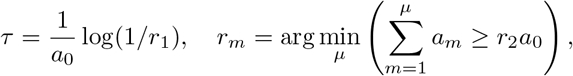

where 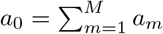 is the total sum of propensities.

In this work, we used the rejection stochastic simulation algorithm (RSSA) [53]. This method estimates the upper and lower bounds on the propensities <u>*a*_*m*_</u>, 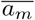, instead of calculating the exact values, *a*_*m*_, and uses the third random number, *r*_3_ *∼ U* (0, 1), for a rejection test to check if the exact value is needed to be computed (see details in Ref. [53]). The propensity values are updated only when necessary, therefore, the algorithm is practical when the propensity computation is time-consuming, e.g., for non-linear process rates parameterised with Michaelis-Menten functions. Additionally, two dependency graph structures are defined to reduce the number of propensity updates and accelerate computations: the first one specifies for each process which variables are affected when the corresponding jumps occur, and the second one specifies for each variable the process indices with propensities dependent on the value of the variable. Note that certain search strategies for the candidate process can also be implemented (e.g., RSSA with composition-rejection search (RSSA-CR) groups the jump processes by their propensity bounds). Alternatively, one can use approximate methods to significantly speed up computations, such as the tau-leaping method [52], or the other hybrid methods that make use of SDE or ODE approximations [53, 54]. In this work, however, we used the exact RSSA method, as the performance of the parallelised code to compute the ensemble of stochastic trajectories was acceptable.

### Software

The following packages in Python (https://www.python.org, version 3.8.8 released 19th February 2021) were used to simulate and analyse the model: Scipy (https://scipy.org/ version 1.8.1, released tbe 20th May 2022) to numerically solve the system of ordinary differential equations, SALib (https://salib.readthedocs.io version 1.4.5) for identification of Sobol sensitivity indices, Matplotlib (https://matplotlib.org/, version 3.5.1 released the 11th December 2021) for visualisations, and Joblib (https://joblib.readthedocs.io, version 1.0.1 released the 9th February 2021) for parallelisation of the ABC rejection algorithm, which allows to infer posterior distributions of model parameters. To perform stochastic simulations, we used the package DifferentialEquations.jl (https://diffeq.sciml.ai/, version 7.4.0) in julia language (https://julialang.org/, version 1.8.1). Codes used to simulate and analyse these models are available in the GitHub repository https://github.com/MacauleyLockeml/SARS-CoV-2-DIP-Model.

## Mathematical model of WT and DIP infection

The variables of the mathematical model characterising the life cycle of SARS-CoV-2 according to Figure 1 are listed in Table 1.

### Cell entry and RNA release

The binding of infectious WT virion to the cellular trans-membrane protein ACE2 allows entry and release of the viral RNA into the host cell. We describe this process by equations specifying the rates of change of free-, receptor-bound, and fused virions, as well as the viral RNA genome in the cytoplasm:

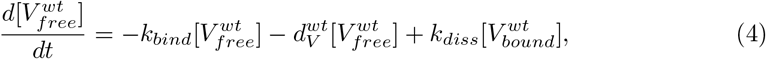

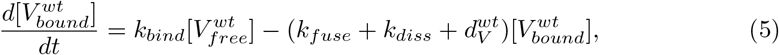

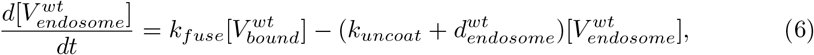

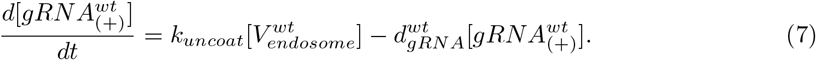

Here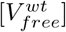 is the number of extra-cellular free infectious virions, 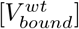, the number of virions bound to ACE2 and activated by TMPRSS2, 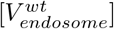, the number of virions in endosomes, and 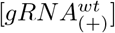, the number of ss-positive sense genomic RNA. A similar set of equations is used to describe the cell entry and RNA release of non-infectious viral defective interfering particles:

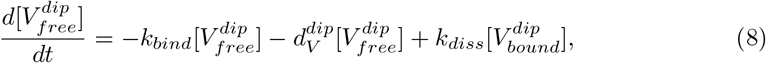

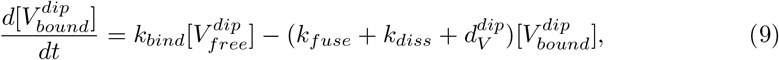

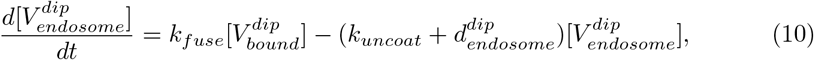

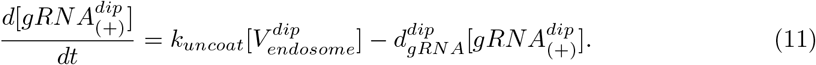

Here 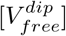 is the number of extra-cellular free DIPs, 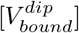, the number of DIPs bound to ACE2 and activated by TMPRSS2, 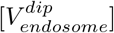, the number of DIPs in endosomes, and 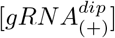, the number of ss-positive sense genomic RNA. DIPs for SARS-CoV-2 would require a functional spike (*S*) protein to successfully bind to ACE2 receptors and mediate cell entry. Consequently, we assume that the rates for *k*_*bind*_, *k*_*diss*_, *k*_*fuse*_, and *k*_*uncoat*_ are the same for both WT virus and DIPs. However, degradation rates related to cell entry will differ between WT and DIPs, since the shorter genome of DIPs might imply a different degradation rate.

### RNA transcription and DIP parasitism

The released WT viral genomic RNA undergoes translation into non-structural viral polyproteins, [*NSP*], which operate to form the viral replication and transcription complex, i.e., the RNA-dependent RNA polymerase (RdRp). The main function of the RdRp replication complex is to generate a negative sense full-length genomic and subgenomic RNAs. As DIPs lack the ability of self-replication, the conditional transcription of DIP RNAs results in competition with WT SARS-CoV-2 for replication proteins [55]. The use of WT virus *trans* elements by DIPs reduces [*NSP*] availability for the transcription of WT viral RNA. The respective sets of equations have different structures, as detailed below. The abundance of non-structural proteins, [*NSP*], the negative sense genomic and subgenomic RNAs, 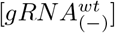, and positive sense genomic and subgenomic RNAs, [*gRNA*^*wt*^], associated with the replication of the infectious virions are described by the following equations:

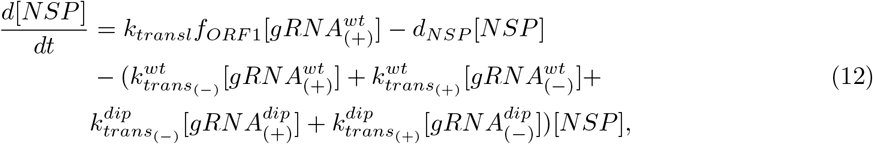

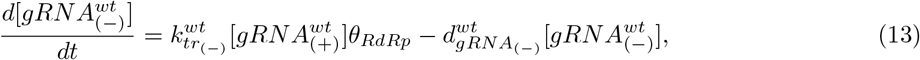

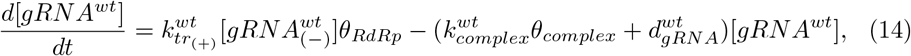

where

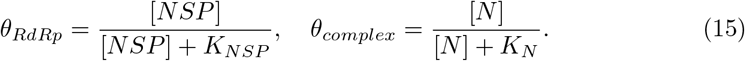

Eq. (12) reflects the fact that non-structural proteins are translated only from the viral genomic RNA of infectious WT virions. Transcription of negative sense viral genomic and subgenomic RNAs described by Eq. (13) and Eq. (14) is regulated by the positive sense viral genomic RNA. The set of equations for transcription of negative sense and positive sense DIP subgenomic RNAs, i.e., 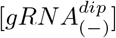, [*gRNA*^*dip*^] are as follows:

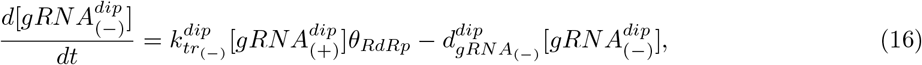

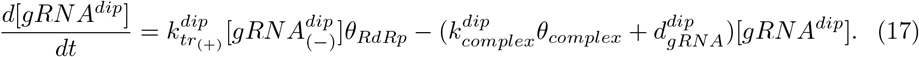

### Translation and competition for nucleocapsid protein and other structural proteins

DIPs compete with WT virions for packaging proteins, e.g., nucleocapsid *N* proteins ([*N*]) [55]. Structural *S*, envelope *E*, and membrane *M* proteins are translated from positive sense subgenomic RNAs in the endoplasmic reticulum (ER) and are considered in the mathematical model as a single population, [*SP*]. Nucleocapsid proteins, on the other hand, are translated in cytosolic ribosomes from positive sense RNAs. Both *SP* and *N* proteins are required for the formation of virus like-particles, WT or DIPs. It can be assumed that 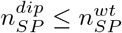 and 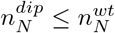, since the shorter DIP genome will require fewer *N* proteins for the formation of the ribonucleocapsid and construction of a viral particle. Translation of *N* and *SP* proteins are described by the following two equations:

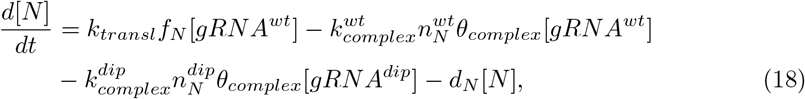

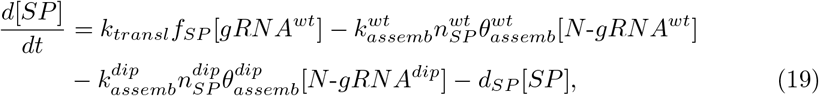

where

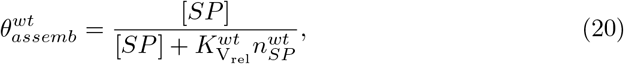

and

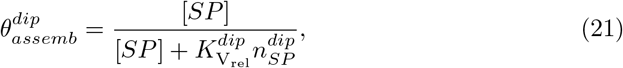

### Assembly and release of WT SARS-CoV-2 and DIPs

New virions are assembled at the endoplasmic reticulum-Golgi compartment, where N-RNA complexes become encapsulated. These assembled virions can then exit an infected cell by exocytosis via the lysosomal pathway, budding, or cell death [56, 57]. There is no competition associated with the release of new infectious and DIP virions, but the viral assembly rates, 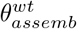 and 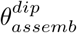, depend on the availability of structural proteins, since DIPs will likely require fewer of them than WT virions. The rates of change of the ribonucleocapsid, assembled and released infectious SARS-CoV-2 and DIPs are described below:

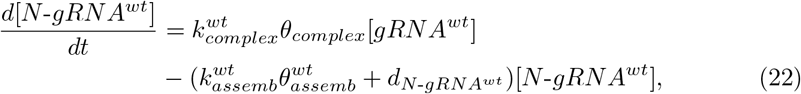

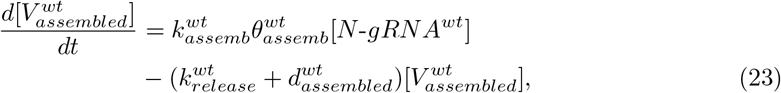

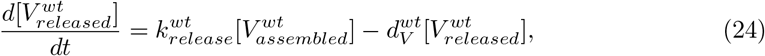

and

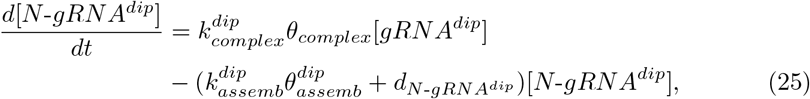

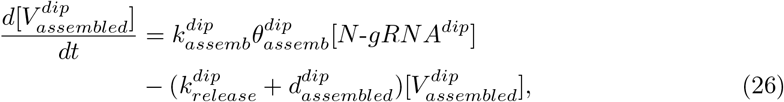

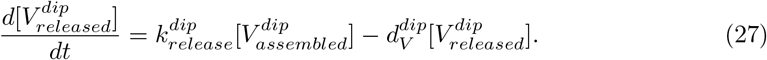

In this study, we wish to explore model behaviour for different initial conditions, 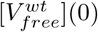 and 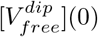, and thus, understand the replication dynamics of WT viral particles in the presence of DIPs, and how the initial dose of WT or DIP particles regulates infection and production kinetics of WT virions. We are also interested in investigating the sensitivities to model parameters of different outputs.

### Stochastic Markov chain model

The deterministic model defined by Eqs. (4)-(27) can be generalised to a stochastic one formulated as a discrete-state continuous-time Markov chain (DSCT MC). The stochastic model allows one to account for integer-valued variables, to obtain probability distributions rather than mean field estimates for the variables of interest, and to compute the probabilities of productive cell infection at low MOI [58]. It is convenient to estimate model parameters for the system of ODEs and then, with a calibrated deterministic system, and a defined Markov chain model, perform stochastic simulations making use of Monte Carlo methods. We follow our previous effort on the stochastic modelling of SARS-CoV-2 [58] and HIV-1 [54] life cycles to formulate and simulate the Markov chain. The Markov chain corresponding to Eqs. (4)-(27) is presented in Table 5. It includes the state transition events and the propensities, *a*_*m*_, for the *m*-th jump process. The propensity *a*_*m*_ defines the probability *p*_*m*_ = *a*_*m*_*dt* that the *m*-th process takes place in the infinitesimal time interval [*t, t* + *dt*). This definition yields exponential distributions for the time between jumps and various Monte Carlo methods can be used to simulate the stochastic trajectories from these distributions [52, 53]. We note that the processes of ribonucleocapsid formation (*m* = 33, 34) and virion assembly (*m* = 37, 38) are formulated as single events, yet involve the simultaneous change of three different variables. In these processes, protein copy numbers are decreased by the corresponding number of protein molecules, *n*_*p*_, needed to form a complex or assemble a pre-virion particle (i.e., by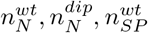, or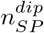, respectively). Alternatively, one can formulate the Markov chain (MC) with three separate processes for each assembly event, in which the protein molecules are decreased by only one molecule with the propensity multiplied by *n*_*p*_ (see Ref. [58] for an example of the extended MC formulation). We have verified that the extended and reduced MCs produce similar statistics. This reduction can be viewed as a weighted sampling strategy used in probability-weighted dynamic Monte Carlo method (PW-DMC) to accelerate computations [53].

**Table 5.** The Markov chain models: individual transitions and their propensities.

| $m$ | Transition | Propensity, $a_m$ |
| --- | --- | --- |
| Entry and RNA release (WT): |  |  |
| 1 | $[V_{free}^{wt}] \rightarrow [V_{free}^{wt}] - 1, [V_{bound}^{wt}] \rightarrow [V_{bound}^{wt}] + 1$ | $k_{bind}[V_{free}^{wt}]$ |
| 2 | $[V_{free}^{wt}] \rightarrow [V_{free}^{wt}] - 1$ | $d_V^{wt}[V_{free}^{wt}]$ |
| 3 | $[V_{free}^{wt}] \rightarrow [V_{free}^{wt}] + 1, [V_{bound}^{wt}] \rightarrow [V_{bound}^{wt}] - 1$ | $k_{diss}[V_{bound}^{wt}]$ |
| 4 | $[V_{bound}^{wt}] \rightarrow [V_{bound}^{wt}] - 1, [V_{endosome}^{wt}] \rightarrow [V_{endosome}^{wt}] + 1$ | $k_{fuse}[V_{bound}^{wt}]$ |
| 5 | $[V_{bound}^{wt}] \rightarrow [V_{bound}^{wt}] - 1$ | $d_V^{wt}[V_{bound}^{wt}]$ |
| 6 | $[V_{endosome}^{wt}] \rightarrow [V_{endosome}^{wt}] - 1, [gRNA_{(+)}^{wt}] \rightarrow [gRNA_{(+)}^{wt}] + 1$ | $k_{uncoat}[V_{endosome}^{wt}]$ |
| 7 | $[V_{endosome}^{wt}] \rightarrow [V_{endosome}^{wt}] - 1$ | $d_{endosome}^{wt}[V_{endosome}^{wt}]$ |
| 8 | $[gRNA_{(+)}^{wt}] \rightarrow [gRNA_{(+)}^{wt}] - 1$ | $d_{gRNA}^{wt}[gRNA_{(+)}^{wt}]$ |
| Entry and RNA release (DIPs): |  |  |
| 9 | $[V_{free}^{dip}] \rightarrow [V_{free}^{dip}] - 1, [V_{bound}^{dip}] \rightarrow [V_{bound}^{dip}] + 1$ | $k_{bind}[V_{free}^{dip}]$ |
| 10 | $[V_{free}^{dip}] \rightarrow [V_{free}^{dip}] - 1$ | $d_V^{dip}[V_{free}^{dip}]$ |
| 11 | $[V_{free}^{dip}] \rightarrow [V_{free}^{dip}] + 1, [V_{bound}^{dip}] \rightarrow [V_{bound}^{dip}] - 1$ | $k_{diss}[V_{bound}^{dip}]$ |
| 12 | $[V_{bound}^{dip}] \rightarrow [V_{bound}^{dip}] - 1, [V_{endosome}^{dip}] \rightarrow [V_{endosome}^{dip}] + 1$ | $k_{fuse}[V_{bound}^{dip}]$ |
| 13 | $[V_{bound}^{dip}] \rightarrow [V_{bound}^{dip}] - 1$ | $d_V^{dip}[V_{bound}^{dip}]$ |
| 14 | $[V_{endosome}^{dip}] \rightarrow [V_{endosome}^{dip}] - 1, [gRNA_{(+)}^{dip}] \rightarrow [gRNA_{(+)}^{dip}] + 1$ | $k_{uncoat}[V_{endosome}^{dip}]$ |
| 15 | $[V_{endosome}^{dip}] \rightarrow [V_{endosome}^{dip}] - 1$ | $d_{endosome}^{dip}[V_{endosome}^{dip}]$ |
| 16 | $[gRNA_{(+)}^{dip}] \rightarrow [gRNA_{(+)}^{dip}] - 1$ | $d_{gRNA}^{dip}[gRNA_{(+)}^{dip}]$ |
| ORF1 translation and competitive viral RNA replication: |  |  |
| 17 | $[NSP] \rightarrow [NSP] + 1$ | $k_{transl}forF1[gRNA_{(+)}^{wt}]$ |
| 18 | $[NSP] \rightarrow [NSP] - 1$ | $d_{NSP}[NSP]$ |
| 19 | $[NSP] \rightarrow [NSP] - 1$ | $k_{trans(-)}^{wt}[gRNA_{(+)}^{wt}][NSP]$ |
| 20 | $[NSP] \rightarrow [NSP] - 1$ | $k_{trans(+)}^{wt}[gRNA_{(-)}^{wt}][NSP]$ |
| 21 | $[NSP] \rightarrow [NSP] - 1$ | $k_{trans(-)}^{dip}[gRNA_{(+)}^{dip}][NSP]$ |
| 22 | $[NSP] \rightarrow [NSP] - 1$ | $k_{trans(+)}^{dip}[gRNA_{(-)}^{dip}][NSP]$ |
| 23 | $[gRNA_{(-)}^{wt}] \rightarrow [gRNA_{(-)}^{wt}] + 1$ | $k_{tr(-)}^{wt}\theta_{RdRp}[gRNA_{(+)}^{wt}]$ |
| 24 | $[gRNA_{(-)}^{wt}] \rightarrow [gRNA_{(-)}^{wt}] - 1$ | $d_{gRNA_{(-)}^{wt}}[gRNA_{(-)}^{wt}]$ |
| 25 | $[gRNA^{wt}] \rightarrow [gRNA^{wt}] + 1$ | $k_{tr(+)}^{wt}\theta_{RdRp}[gRNA_{(-)}^{wt}]$ |
| 26 | $[gRNA^{wt}] \rightarrow [gRNA^{wt}] - 1$ | $d_{gRNA}^{wt}[gRNA^{wt}]$ |
| 27 | $[gRNA_{(-)}^{dip}] \rightarrow [gRNA_{(-)}^{dip}] + 1$ | $k_{tr(-)}^{dip}\theta_{RdRp}[gRNA_{(+)}^{dip}]$ |
| 28 | $[gRNA_{(-)}^{dip}] \rightarrow [gRNA_{(-)}^{dip}] - 1$ | $d_{gRNA_{(-)}^{dip}}[gRNA_{(-)}^{dip}]$ |
| 29 | $[gRNA^{dip}] \rightarrow [gRNA^{dip}] + 1$ | $k_{tr(+)}^{dip}\theta_{RdRp}[gRNA_{(-)}^{dip}]$ |
| 30 | $[gRNA^{dip}] \rightarrow [gRNA^{dip}] - 1$ | $d_{gRNA}^{dip}[gRNA^{dip}]$ |
| Translation and ribonucleocapsid formation: |  |  |
| 31 | $[N] \rightarrow [N] + 1$ | $k_{transl}f_N[gRNA^{wt}]$ |
| 32 | $[N] \rightarrow [N] - 1$ | $d_N[N]$ |
| 33 | $[gRNA^{wt}] \rightarrow [gRNA^{wt}] - 1, [N] \rightarrow [N] - n_N^{wt},$<br>$[N-gRNA^{wt}] \rightarrow [N-gRNA^{wt}] + 1$ | $k_{complex}^{wt}\theta_{complex}[gRNA^{wt}]$ |
| 34 | $[gRNA^{dip}] \rightarrow [gRNA^{dip}] - 1, [N] \rightarrow [N] - n_N^{dip},$<br>$[N-gRNA^{dip}] \rightarrow [N-gRNA^{dip}] + 1$ | $k_{complex}^{dip}\theta_{complex}[gRNA^{dip}]$ |
| 35 | $[SP] \rightarrow [SP] + 1$ | $k_{transl}f_{SP}[gRNA^{wt}]$ |
| 36 | $[SP] \rightarrow [SP] - 1$ | $d_{SP}[SP]$ |
| Assembly and release: |  |  |
| 37 | $[N-gRNA^{wt}] \rightarrow [N-gRNA^{wt}] - 1, [SP] \rightarrow [SP] - n_{SP}^{wt},$<br>$[V_{assembled}^{wt}] \rightarrow [V_{assembled}^{wt}] + 1$ | $k_{assemb}^{wt}\theta_{assemb}^{wt}[N-gRNA^{wt}]$ |
| 38 | $[N-gRNA^{dip}] \rightarrow [N-gRNA^{dip}] - 1, [SP] \rightarrow [SP] - n_{SP}^{dip},$<br>$[V_{assembled}^{dip}] \rightarrow [V_{assembled}^{dip}] + 1$ | $k_{assemb}^{dip}\theta_{assemb}^{dip}[N-gRNA^{dip}]$ |
| 39 | $[N-gRNA^{wt}] \rightarrow [N-gRNA^{wt}] - 1$ | $d_{N-gRNA}^{wt}[N-gRNA^{wt}]$ |
| 40 | $[N-gRNA^{dip}] \rightarrow [N-gRNA^{dip}] - 1$ | $d_{N-gRNA}^{dip}[N-gRNA^{dip}]$ |
| 41 | $[V_{assembled}^{wt}] \rightarrow [V_{assembled}^{wt}] - 1, [V_{released}^{wt}] \rightarrow [V_{released}^{wt}] + 1$ | $k_{release}^{wt}[V_{assembled}^{wt}]$ |
| 42 | $[V_{assembled}^{wt}] \rightarrow [V_{assembled}^{wt}] - 1$ | $d_{assembled}^{wt}[V_{assembled}^{wt}]$ |
| 43 | $[V_{released}^{wt}] \rightarrow [V_{released}^{wt}] - 1$ | $d_V^{wt}[V_{released}^{wt}]$ |
| 44 | $[V_{assembled}^{dip}] \rightarrow [V_{assembled}^{dip}] - 1, [V_{released}^{dip}] \rightarrow [V_{released}^{dip}] + 1$ | $k_{release}^{dip}[V_{assembled}^{dip}]$ |
| 45 | $[V_{assembled}^{dip}] \rightarrow [V_{assembled}^{dip}] - 1$ | $d_{assembled}^{dip}[V_{assembled}^{dip}]$ |
| 46 | $[V_{released}^{dip}] \rightarrow [V_{released}^{dip}] - 1$ | $d_V^{dip}[V_{released}^{dip}]$ |

## Results

### Sensitivity analysis

We now evaluate how model outputs change with parameter values. To that end a Sobol global sensitivity analysis was performed on four different model outputs. We first considered the variability of WT genomic RNA, [*gRNA*^*wt*^], and DIP genomic RNA, [*gRNA*^*dip*^], as a result of modifying parameter values within a set range summarised in Table 2 and Table 4. Secondly, we investigated how parameter variability affects the release kinetics of both WT 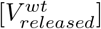 and DIP 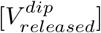 particles 48 hours post-infection. Understanding which parameters cause the most variability in our model will allow us to calibrate it with careful consideration to minimise output perturbations.

Figure 2 illustrates the first and total order sensitivities for WT genomic RNA, [*gRNA*^*wt*^], and DIP genomic RNA, [*gRNA*^*dip*^], as outputs of the proposed model. For [*gRNA*^*dip*^], the parameter 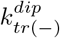 was identified as generating the largest variation. 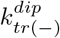 is associated with the transcription of negative sense RNAs for DIPs, and thus, is essential in the formation of new positive sense genomic and subgenomic RNAs. The rate 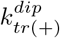 was also identified as a high sensitivity parameter, since it is associated with the transcription of positive sense RNAs. Consequently, 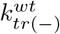 was the second most important parameter in minimising variation in model output for [*gRNA*^*wt*^], following the same reasoning as for DIP positive sense genomic RNA.

**Fig 2.**
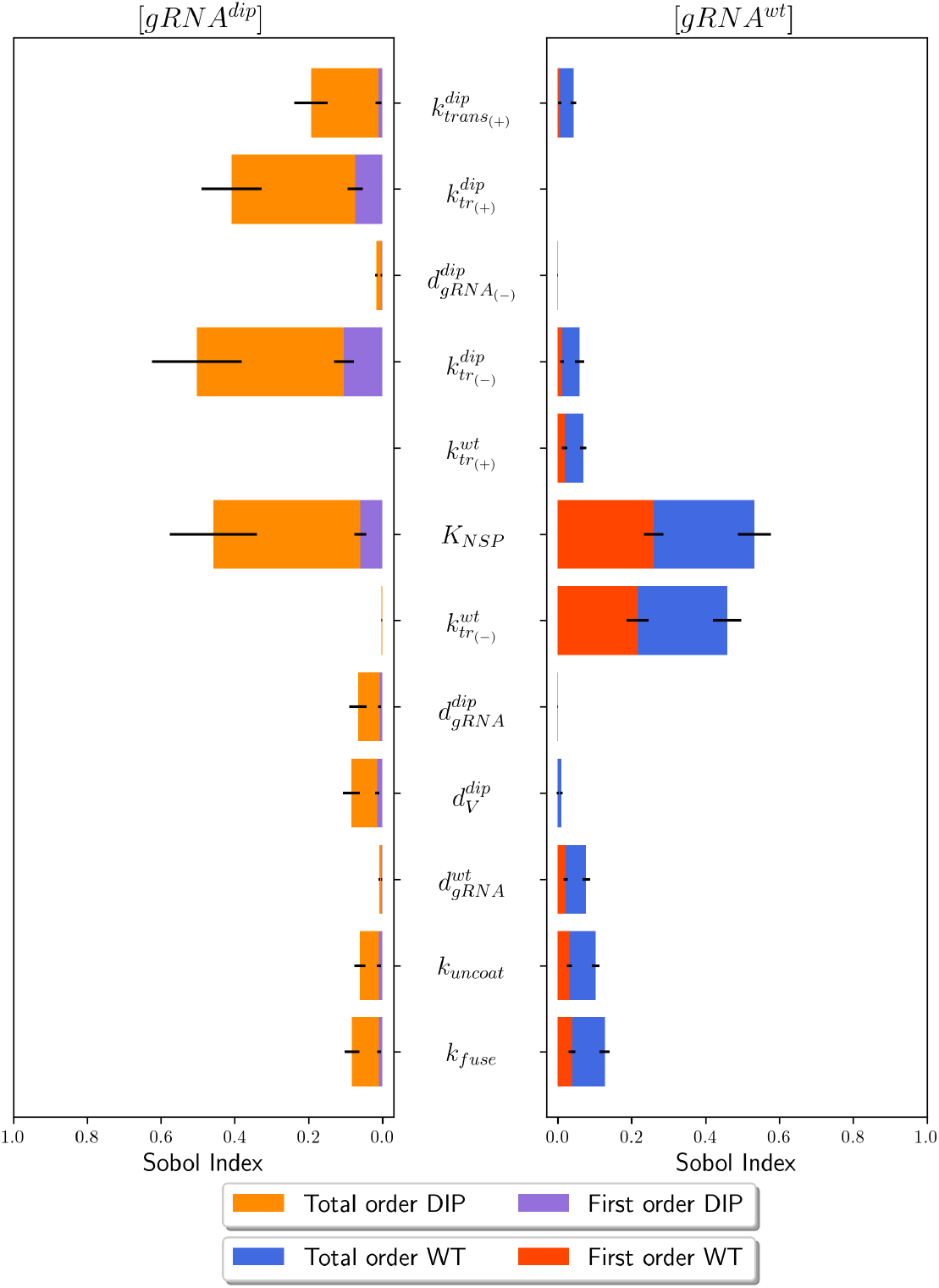
Sensitivity analysis for positive sense genomic RNA. First and total order sensitivities from 10^4^ samples with 95% confidence interval (black line). **(Left:)** sensitivities for the variable [*gRNA*^*dip*^](48). **(Right:)** sensitivities for the variable [*gRNA*^*wt*^](48). Initial conditions used for sensitivity analysis were 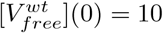 and 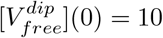.

**Fig 3.**
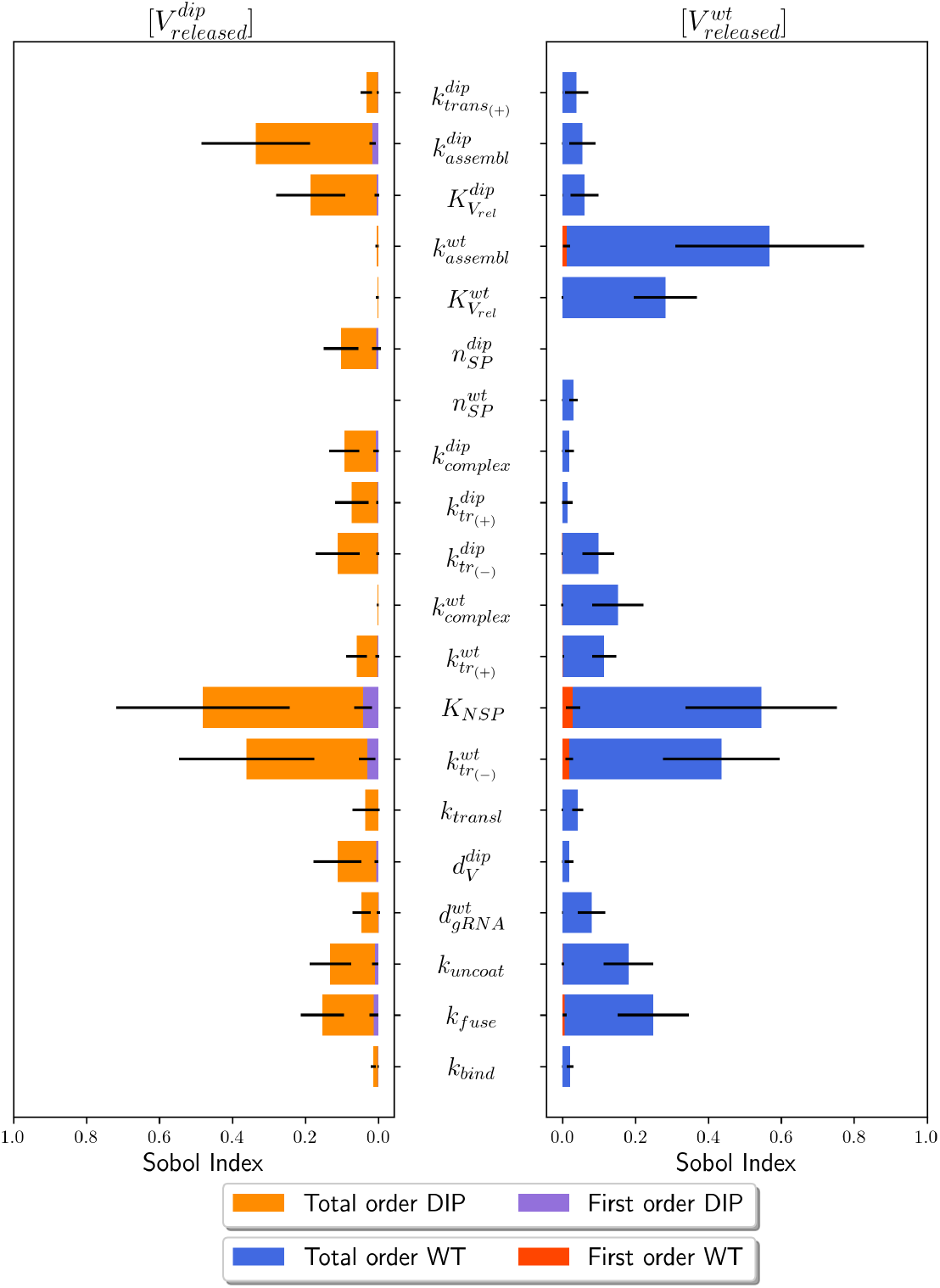
Sensitivity analysis for released particles. First and total order sensitivities from 10^4^ samples with 95% confidence interval (black line). **(Left:)** sensitivities for the variable 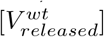 (48). **(Right:)** sensitivities for the variable 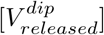 (48). Initial conditions of 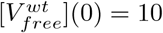 and 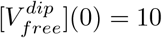 were used for sensitivity analysis.

A parameter that was of great importance, and not only caused large variation in model outputs of [*gRNA*] for WT or DIPs, but also [*V*_*released*_], was the threshold parameter of non-structural proteins, *K*_*NSP*_. *K*_*NSP*_ causes the most variation for 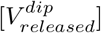 and [*gRNA*^*wt*^] compared to any other parameter, and for 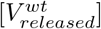 and [*gRNA*^*dip*^] it is the second most important parameter. *K*_*NSP*_ is associated with the transcription of both negative and positive sense genomic RNAs, and changes in the value of this parameter will modify the number of WT virions and DIPs released. 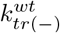 was identified as an important parameter to minimise variation in the release of both WT 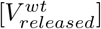 and DIPs 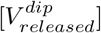. Consequently, transcription of negative sense WT genomic RNAs is an essential first step in producing positive stranded gRNA, which is then translated to form structural proteins *S, M*, and *E* ([*SP*]), as well as nucleocapsid proteins ([*N*]) which are required to form new viral particles. Parameters associated with WT virion or DIP assembly are also important to monitor to reduce variation in model outputs. Several of the parameters identified by the Sobol sensitivity analysis have been previously estimated in Ref. [21] and are summarised in Table 2. Other parameters required estimation and these are listed in Table 4.

### Parameter calibration

In our extension of the model proposed by Grebennikov *et al*. in Ref. [21], we introduced several parameters which have not been previously quantified. To estimate their values, we performed Bayesian parameter calibration. Since experimental data sets on co-infection with DIPs is limited, we aimed to achieve the fold reduction experimentally quantified by Chaturvedi *et al*. in Ref. [15]. We made use of an ABC rejection algorithm with 10^6^ sample sets. As previously mentioned, since a choice of *ε* is hard to determine, we instead took the 0.1% of parameter sets which minimise the Euclidean distance. We sampled the exponent of the search ranges shown in Table 4. As a result, our sample size provided a large coverage of parameter space. We compared the fold log reduction between the reference solution of a model without DIPs and the one with DIPS to the data in Table 3, and Figure 4 illustrates the model output where we used the median values from the accepted 0.1% sample sets. From these median values we obtained a fold change of 1.08 (two d.p.) at 24 hours post-infection and 1.14 (two d.p.) at 48 hours post-infection, compared to the reference solution [21]. Posterior histograms in Figure S1 showed that with the data set and the mathematical model, Bayesian inference has led to poor learning for all but one of the newly introduced parameters. Posterior distributions are still extremely wide, with 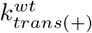 being the only parameter with a narrow posterior distribution. This was due to lack of longitudinal data to compare modelled DIP replication dynamics with.

**Fig 4.**
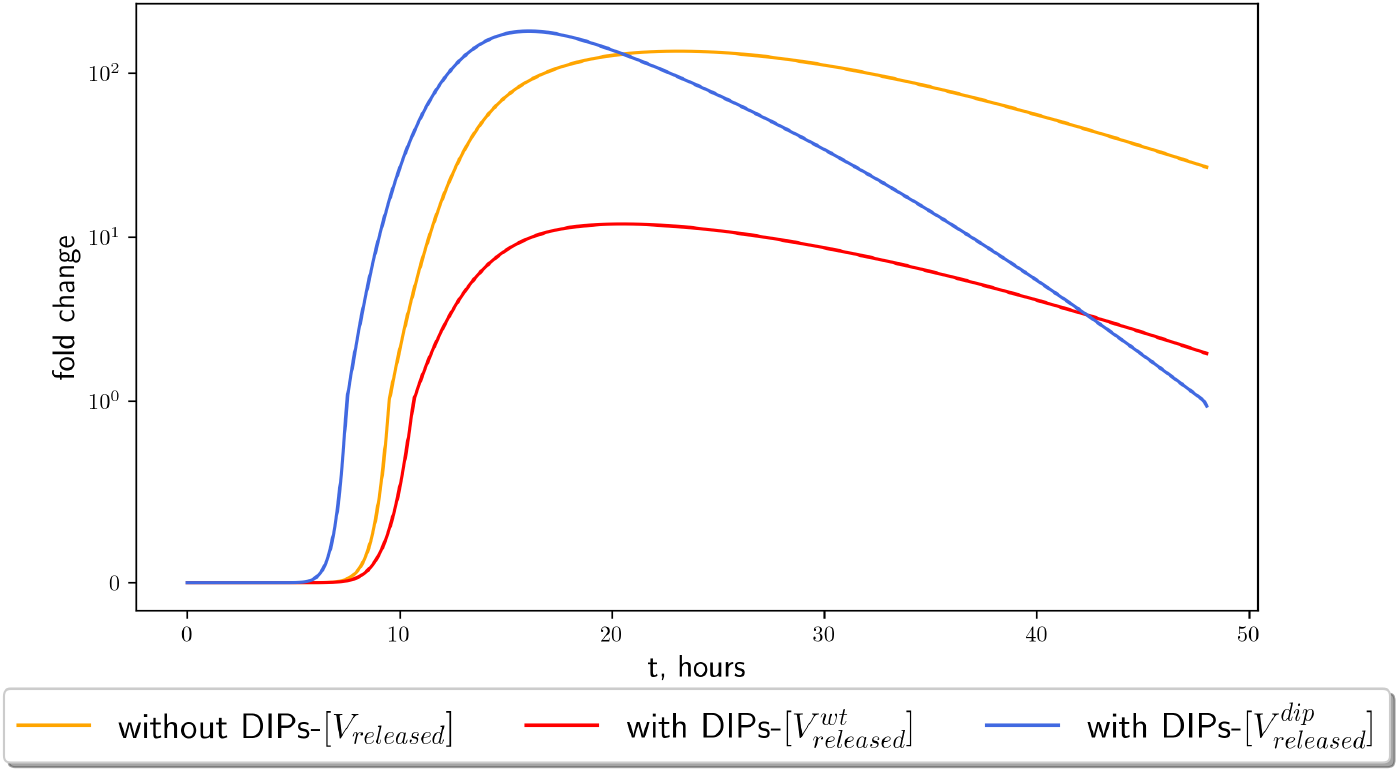
Time evolution of viral particle release for the parameter values estimated with the ABC method. Viral particle release kinetics predicted by the model with initial conditions 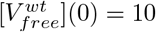 and 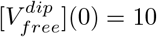 48 hours post-infection after model calibration using ABC rejection and data from Chaturvedi *et al*. [15]. Median parameter values summarised in Table 4 were used for previously unknown parameter values. **(Yellow line:)** shows the reference solution to a model where DIPs are not considered in the replication dynamics. **(Red line:)** illustrates the production of WT virions 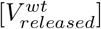 with DIPs **(blue line)** 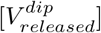.

Figure 5 illustrates the time evolution for each variable in Table 1 given the median values found via ABC rejection. From the upper panels of Figure 5 we examined that the entry kinetics of WT virus into the cell are similar to those of the reference solution. DIPs, however, enter the cell at a faster rate than WT virions. It is important to remember that we assumed there are sufficient ACE2 receptors mediating viral entry; thus, there is no competition between WT and DIP for receptor binding. The number of non-structural proteins is greatly reduced (Figure 5 middle left panel), peaking at 7 hours with *≈* 20 molecules as opposed to the reference solution, which peaks at roughly 13 hours with *≈* 40 molecules. The production of 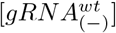 halves and peaks earlier in the time course, with a greater number of DIP negative sense genomic RNA than WT. Consequently, we then saw an approximate fold reduction of positive sense genomic RNA, ribonucleocapsid proteins, assembled and released WT viral particles.

**Fig 5.**
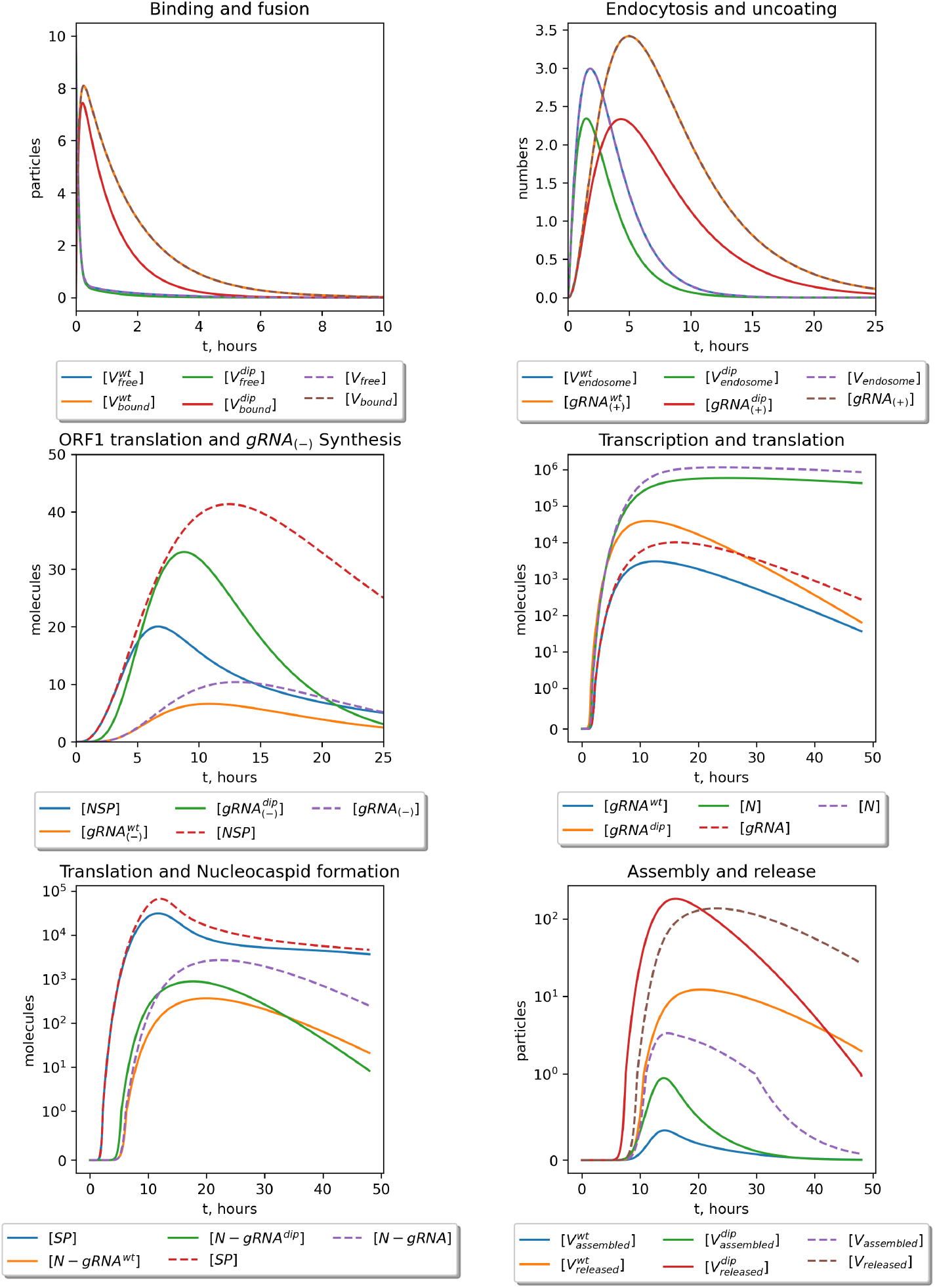
Deterministic model outputs. Time-dependent state variables of the mathematical model for the life cycle of SARS-CoV-2, including wild-type virions and defective interfering particles with initial conditions 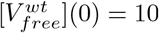 and 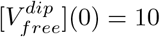 over a 48 hour time course. **(Upper left:)** free WT or DIP virions bind and fuse to the cell ACE2 receptors, and **(upper right:)** virions entering endosomes and uncoating of viral positive sense genomic RNA. **(Middle left:)** transcription and translation to form a negative sense genome and ORF1 to form non-structural proteins (*NSP* s), which is then followed by **(middle right:)** the production of new positive sense genomic RNAs and translation of *N* protein. **(Bottom left:)** translation of structural proteins and formation of ribonucleocapsid molecules, which lead to **(bottom right:)** the assembly and release of new virions, both WT and DIP. **(Dashed lines:)** represent the reference model solution proposed in Ref. [21].

### Stochastic model results

Figure 6 shows the kinetics of the stochastic model variables for initial doses of wild-type virus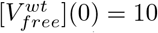 and DIPs 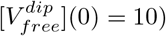. The figure illustrates parametric (mean values) and non-parametric (medians, inter-quartile ranges) statistics computed on an ensemble of 10^6^ trajectories. Additionally, the histograms of the simulated variable values at particular time points can be produced from the ensemble for analysis (Figure S2). The means and medians follow approximately the deterministic model outputs, while the inter-quartile ranges estimate the uncertainty of the simulations due to stochastic effects caused by the discrete nature of the model variables. These stochastic effects are more prominent for variables which are present in a cell in small numbers, and the assumption that their mean values can be approximated by the deterministic model may not be satisfied. In particular, the deterministic model predicts that an infection is productive for every positive initial dose of 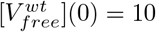, while the stochastic trajectories can become extinct due to stochasticity. Figure 7 illustrates the probability of productive infection as a function of the initial WT virion (MOI) and DIP doses. As can be seen in Figure 7 (left panel), the probability of a productive infection tends to one as the initial dose of the WT virus hits 20 viral particles. However, the probability is affected by the initial dose of DIP particles (Figure 7, right panel), with this probability being reduced linearly as the dose increases.

**Fig 6.**
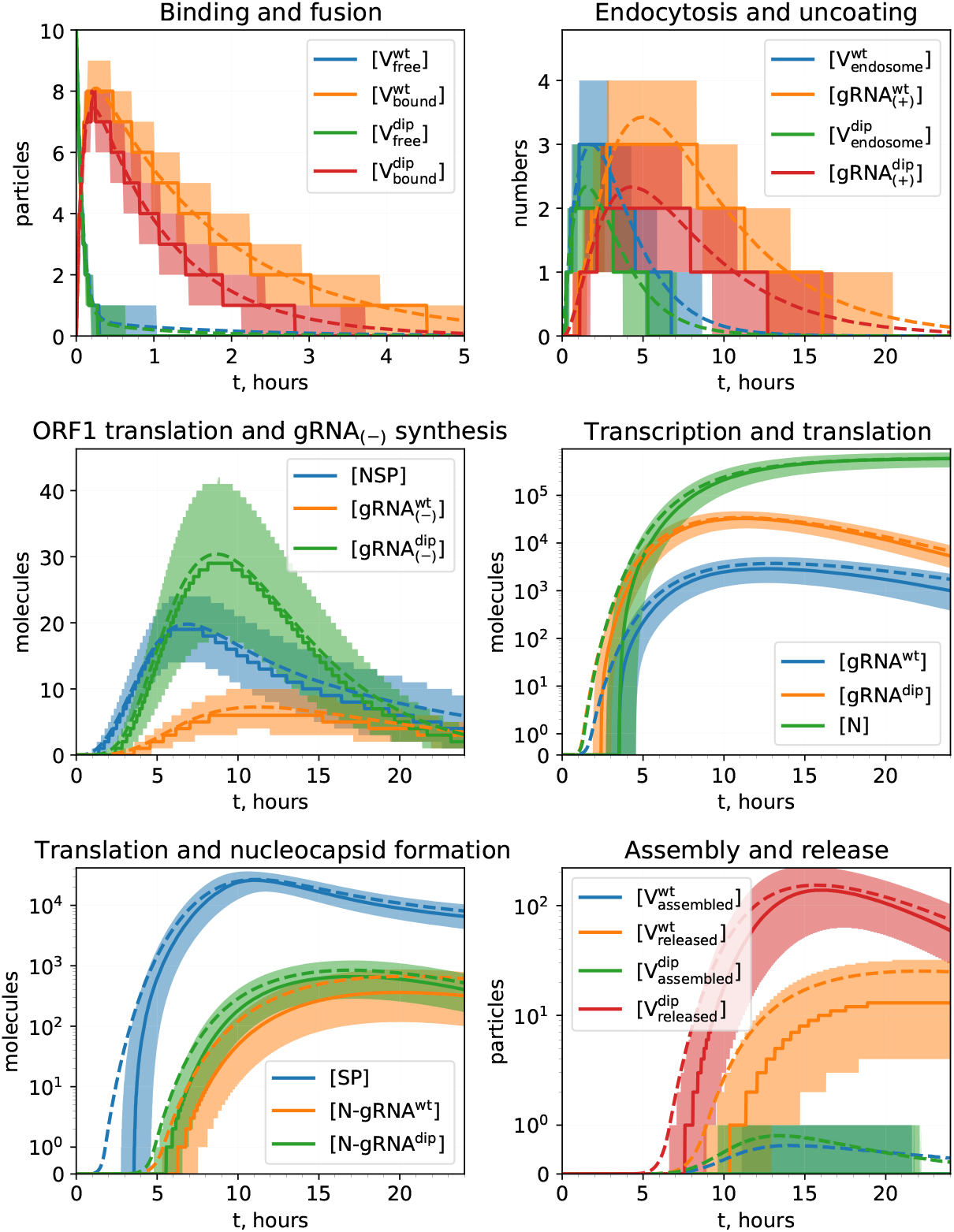
Stochastic model outputs. Statistics of time-dependent state variables of the stochastic model with initial conditions 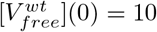 and 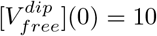 over a 24 hour time course based on an ensemble of 10^6^ simulated trajectories. **Solid lines:** medians, **dashed lines:** mean values, **filled areas:** inter-quartile ranges.

**Fig 7.**
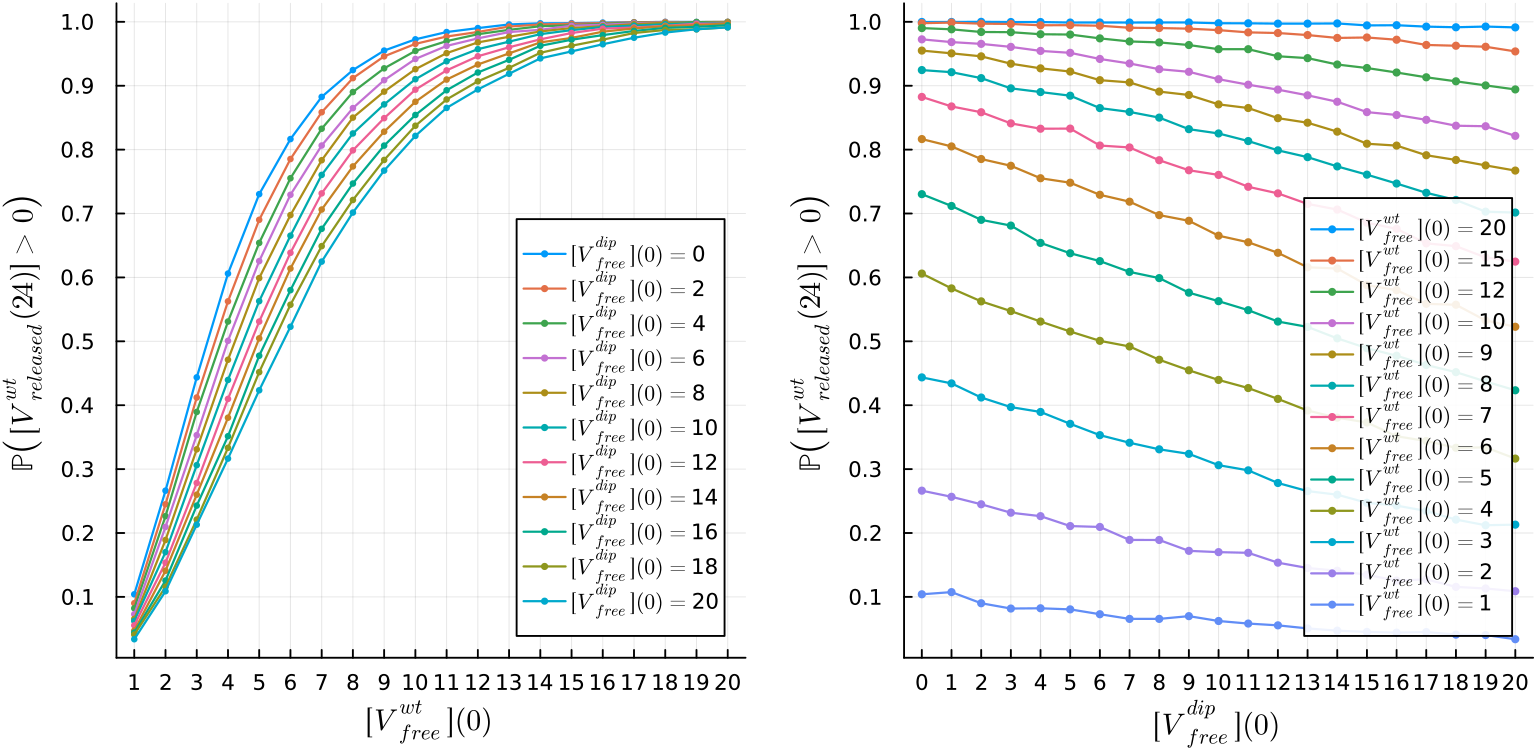
Probability of productive infection. Probability of a productive infection for varying initial doses of both WT 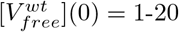 and DIPs 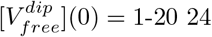 hours post-infection. **Left:** dependence of probability on 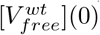 for various 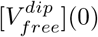, **right:** dependence of probability on 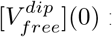 for various 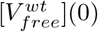.

The mean values of WT virion and DIP production 24 hours post-infection closely follow the outputs predicted by the deterministic model. However, a part of the trajectories simulated with the stochastic model become extinct. The probability of productive infection as a function of WT *MOI* and *DIP*_0_ is shown in Figure 7. When this probability is close to one, a change in *DIP*_0_ does not significantly modify it. For every WT *MOI*, an increase in DIP dose reduces this probability linearly. Figure S5 shows the dependence of this linear decay in probability, *β*_*wt*_, on WT *MOI*.

### Dose response analysis

We examined the release kinetics, i.e., the abundance of WT virions compared to DIPs, as a function of the initial doses 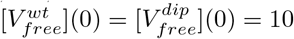. However, one can expect that initial infection doses might vary from cell to cell. Therefore, we now examined the release kinetics of WT virions under different initial conditions. Figure 8 illustrates the total number of WT virions (**left**) and DIPs (**right**) released with initial conditions 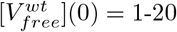 and 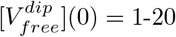 over a 24 hour time period. As can be seen from Figure 8, a low dose of DIPs (MOI = 1) with a high dose of WT virus (MOI = 20) results in an approximately 21% reduction of the WT particles released during DIP co-infection. Furthermore, as we decreased the initial number of WT virions while DIPs remained at an MOI = 1, we observed a continued decrease in WT virus released during co-infection compared to the single infection case (no DIPs). As the dose of DIPs was increased, the total number of WT virions released rapidly decreased, and at MOI=10 for WT and DIP MOI = 4 WT particles only account for approximately 30% of particles released. These deterministic results were consistent with median estimates from the stochastic model presented in Figure S3 (upper panel), while the mean estimates (Figure S3, lower panel) showed marginally higher release in WT virus and lower release of DIPs. Additionally, for high doses of WT virus, a productive infection is almost guaranteed (Figure 7), but as shown in Figure S3, even if an infection is guaranteed the overall number of WT, and hence infectious particles, is reduced.

**Fig 8.**
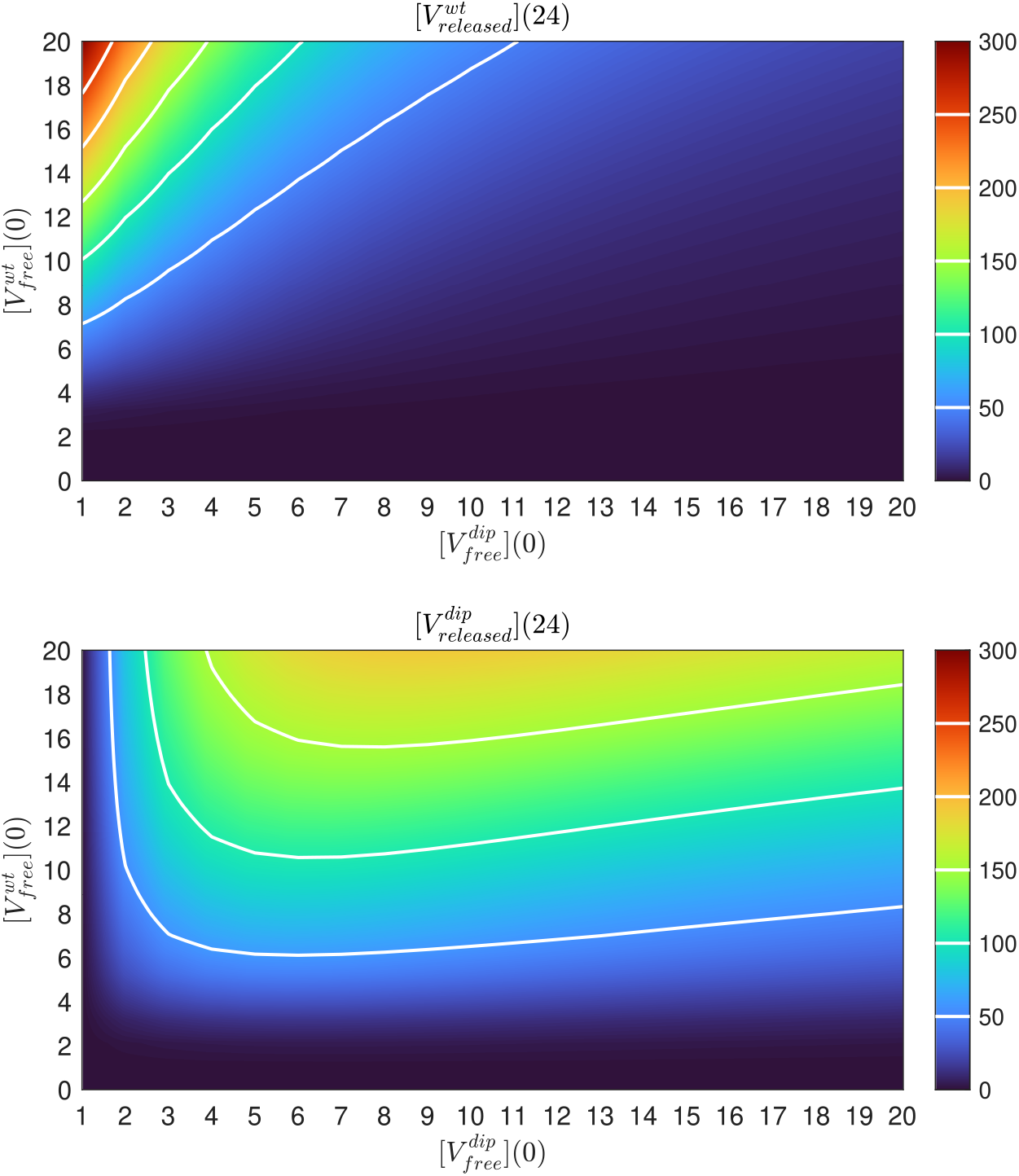
Effects of varying initial dose on viral particle release. **Top:** Total WT virions released over the 24 hours post-infection for varying initial conditions of free WT virions 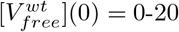 and free DIPs 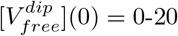 from the deterministic model. **Bottom:** Total DIP particles released for varying initial doses. The isolines shown on the heatmaps as white lines coincide with the corresponding ticks in the colorbars.

Figure 9 shows viral particle release kinetics predicted by the deterministic model with fixed initial conditions for 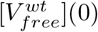 ranging from 3 to 20 virions and varying initial conditions for DIPs 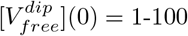. DIP release peaks at a MOI = 6 and then begins to decrease as the dose increases. An increase in dose continues to have an effect on the release of WT virions, so that for a MOI = 40 total WT virion production is *<* 30 virions released over the 24 hour time period considered. This highlights the ability of DIPs to compete (with an advantage) for replication resources with WT virions. Consequently, if the dose is high enough, DIPs sequester so many intra-cellular resources that WT production is significantly reduced. Finally, the non-linear effects of DIP MOI on WT virion and DIP production per cell suggest that there might be optimal dosing of DIPs when used as a therapeutic agent. The maximum effect can potentially be achieved at around 5 to 10 DIPs per cell as this would maximise the number of new DIPs produced by the infected cells, and these, in turn, will reduce the WT virion production in other infected bystander cells.

**Fig 9.**
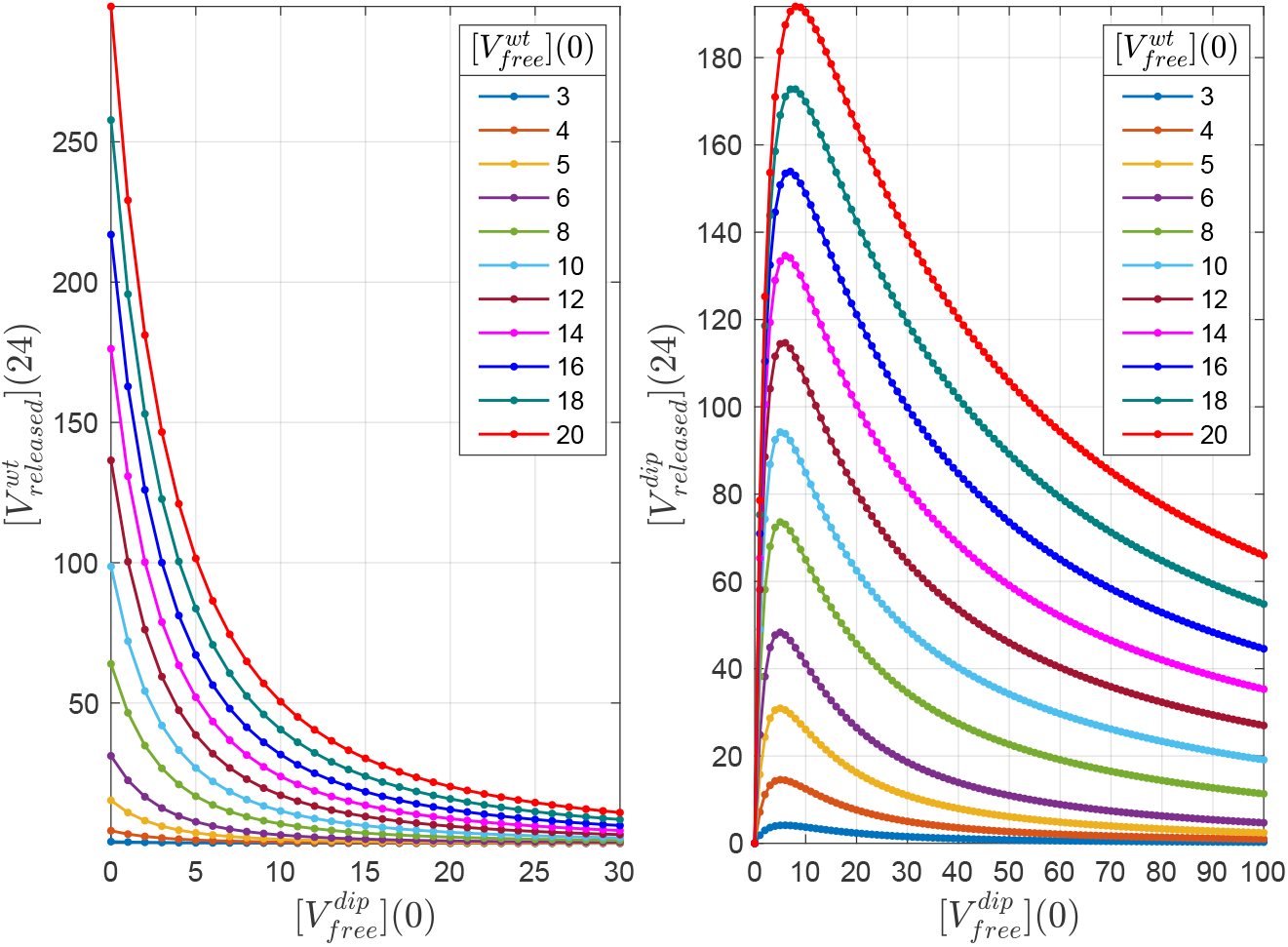
Total WT virions and DIPs released for increased initial doses of DIPs. Viral particle release kinetics predicted by the model with fixed initial conditions for 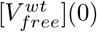 ranging from 3 to 20 virions, as a function of varying DIP doses 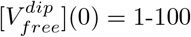.

### DIP dose effect on WT virion production

Given the predicted three-dimensional curves of model outputs as function of initial doses presented as heatmaps in Figure 8, we asked if the production of WT virions 24 hours post-infection 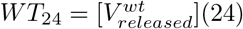 as function of initial doses 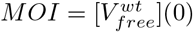 and 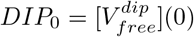 can be approximated by a compact analytic expression. Figure 9 shows that *WT*_24_ as a function of *DIP*_0_, for a fixed *MOI*, exhibits a decay that is slower than exponential (which would be displayed as a straight line on a logarithmic scale). Therefore, we used several analytical expressions with slower than exponential decay to fit the deterministic model predictions for WT virion production for fixed *MOI* = 10. These include: (a) a Gompertz curve, and the probability density functions (p.d.f.) of (b) power-law, (c) Weibull, (d) Cauchy, (e) Burr, (f) Lomax, and (g) generalised Pareto heavy-tailed distributions. The error that was minimized is the sum of squares between the WT virion production, *WT*_24_, predicted by an analytic expression and predicted by the deterministic model for each *DIP*_0_ ranging from 1 to 100. The generalised Pareto distribution (with a location parameter equal to zero) was chosen as the optimal analytic expression making use of the Akaike information criterion. The parameters of the generalised Pareto distribution *ξ* (shape) and *σ* (scale) can be fitted for different values of *MOI*, thus giving the functions *ξ*(*MOI*) and *σ*(*MOI*). The overall parameterisation is the following:

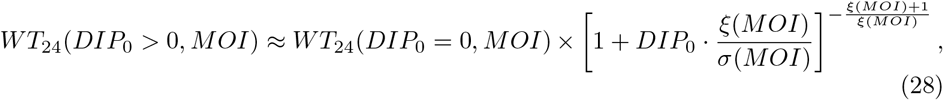

where *WT*_24_(*DIP*_0_ = 0, *MOI*) is the number of released WT virions 24 hours post-infection with zero DIP initial dose for a given WT *MOI*.

Figure S4 shows the fit of a generalised Pareto function (28) for *MOI* = 10 and the dependence of parameters *ξ* and *σ* on *MOI*. One can see that the fit follows the data closely for suitable numbers of produced WT virions (*WT*_24_ *>* 1) and has some small discrepancies for *WT*_24_ *<* 1 at large DIP doses *DIP*_0_ *>* 40. The dependences of parameters *ξ*(*MOI*) and 1*/σ*(*MOI*) exhibit non-linear patterns. They can be approximated with a Hill function and a Dagum distribution p.d.f., respectively. However, when these analytic approximations are substituted in (28), the overall fit of (28) behaves approximately as an exponential decay function (data not shown). Therefore, one should use the computed estimates of the parameters *ξ* and *σ* for every *MOI*, or approximate them with a higher degree polynomial that would follow the estimates closely, e.g., with a 30-degree Chebyshev polynomial as shown in Figure S4. Overall, the relative error of the fit (weighted residual sum of squares) of the closed-form expression (28) reaches its peak for *MOI ≈* 6, in the same region where the parameters *ξ* and *σ* shown a non-linear dependence on *MOI*. The root-mean-square deviation normalised to *WT*_24_(*DIP*_0_ = 0, *MOI*), the maximum value of produced WT virions for each MOI, shows a similar increase near *MOI ≈* 10, as well as a later increase for large MOIs. This can be explained since the discrepancy in the tail of a generalised Pareto distribution corresponds to larger numbers of *WT*_24_ with an increase of *MOI*. In summary, we have provided a closed-form expression, (28), as a prediction of the effect of DIPs on productive cell infection, i.e., the expected mean number of WT virions produced in a productive infection scenario for a range of relevant MOIs.

## Discussion

SARS-CoV-2 still presents a real threat to human health as a result of several compounding factors: emergence of new strains due to mutation, waning immunity amongst the vaccinated, and un-vaccinated individuals (for medical reasons or personal choice). Therefore, it is still important to investigate new treatment options, especially those that could be implemented early after infection, to alleviate pressure on healthcare systems. One such potential therapy is defective interfering particles. DIPs are virus-like particles with shorter genomes that require a wild-type (WT) virus to replicate. In this paper, we investigated the intra-cellular replication kinetics of WT virus in the presence of DIPs, making use of a mathematical model. To this end, we extended the model proposed by Grebennikov *et al*. in Ref. [21], which focused on the intra-cellular replication kinetics of SARS-CoV-2, to include co-infection with defective interfering particles, given their therapeutic potential [59, 60]. In particular, we investigated the ability of DIPs to reduce WT viral load by competing for resources required to replicate or encapsulate the viral genome to form new virions. Since DIP genomes lack key fragments, they need a “helper” virus, which encodes non-structural and structural proteins, for their replication. There is evidence of DIPs leading to cause a reduction in viral titres for several viruses including: influenza A, dengue fever and SARS-CoV-2 [14, 15, 61]. With the emergence of new SARS-CoV-2 strains, the effectiveness of a DIP particle (derived from a particular viral strain) against novel ones remains to be investigated.

Mathematical models of WT virus and DIP co-infection have been investigated at the within host-level and either consider: a standard infection model with target, eclipse phase and infected cells or include different localised areas of infection, such as the upper and lower respiratory tract [15, 19]. There are, however, no models (to the best of our knowledge) that examine the intra-cellular replication kinetics of SARS-CoV-2 in the presence of DIPs. Our aim was to assess the hypothesis that DIPs lead to a reduction not only in the number of released WT virions but also, negatively impact the transcription of positive sense genomic RNAs. Additionally, we investigated the effects of initial infection dose (WT and DIP) in the release of both new WT virions and DIPs. Since experimental data sets are extremely limited, it is important to note that the parameter values obtained in this manuscript are based on the data set used [15], and may not be globally identifiable. By globally identifiable we mean the identification of a unique parameter value from a data set.

The extension of the model presented by Grebennikov *et al*. in Ref. [21] required new parameters to account for the kinetics of DIPs. Therefore, it was necessary to investigate the sensitivities of all model parameters. In particular, we made use of the Sobol sensitivity analysis to understand how variation in parameter values affects four different model outputs: [*gRNA*^*wt*^], [*gRNA*^*dip*^], 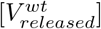, and 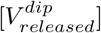. We found several parameters that have an effect on all four model outputs: *K*_*NSP*_, the threshold number of non-structural proteins, 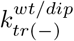, transcription rates of negative sense genomic RNA for WT virus and DIPs, respectively, and 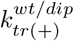, the transcription rates for positive sense genomic RNA. The rates associated with cell entry, *k*_*fuse*_ and *k*_*uncoat*_, also lead to some variation in model outputs. Finally, if we examine as output WT and DIP release, we find their associated assembly rates, 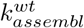 and 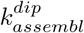, as the most sensitive parameters.

DIPs have potential as therapeutics, thus, it is important to explore how initial infection doses of WT and DIP alter the release of WT virus, to inform a treatment plan. We show that even a low MOI= 1 of DIPs can cause a reduction of approximately 50% in released WT virus compared to an infection in the absence of DIPs, with further reduction in released WT up to 10-fold for increasing MOI_*wt*_ and MOI_*dip*_. Figure 8 illustrates how increasing the dose of DIPs leads to a reduction in the fraction of released WT virions, in relation to the initial WT infection dose. These trends are consistent with the results from the stochastic model also developed in this paper (Figure S3). The doses of both WT virus and DIPs also had an effect on the probability of a productive infection, which decreased with increased doses of DIPs, but is almost certain for high doses of WT virus. We also investigated the effect of initial MOI of DIPs given a fixed dose of WT virus (MOI=10) on viral particle release. Our results show that while DIP release peaks at an initial DIP dose of MOI=5, the release of WT virions decreases in a dose-dependent manner. Furthermore, by an initial DIP dose of MOI=40, WT virion release is effectively inhibited.

The deterministic and stochastic models we presented are a good first approximation to the kinetics of WT and DIP co-infection. Yet, there are a number of biological processes which have not been considered. First and foremost, we omitted the anti-viral response of the cell. While we need not consider the adaptive immune response since our time interval is 48 hours, the innate immune response would play a pivotal role [62, 63]. A family of cytosolic receptors, known as pattern recognition receptors (PRR), exists that detect viral RNAs to induce the production of type I interferons. Type I interferons (or viral IFNs), which are secreted by infected cells, include IFN-*α*, IFN-*β*, IFN-*ω* and IFN-*τ*. These molecules are associated with activation of anti-viral cell states, which in turn lead to inhibition of viral replication and eventual viral clearance [64]. Furthermore, innate immune responses have been shown to be induced by DIP binding to PRRs, providing additional stimuli and magnifying the anti-viral cellular response [59]. As a consequence, it would, therefore be ideal to extend the proposed model to consider the role of an innate immune response. Another limitation of our model is that for WT virions, we do not distinguish between infectious and non-infectious particles. This would be important to understand the potential infectivity of the viral particles released. We also fail to characterize the natural generation of DIPs during the WT replication cycle (which is inherently characterised by mutations). This process would contribute to the release of other defective interfering particles, and would potentially reduce the number of WT virions released. However, a complete calibration of such a model would require a data set not currently at hand.

To conclude, we believe the model we have proposed shows the potential benefits of DIPs as a therapeutic tool to reduce WT virus production. We also have shown that even low doses of these particles can have a positive effect on limiting WT virus production and reducing the probability of a successful infection. This reduction continues, in a dose dependent manner, to greatly reduce WT virus production. Future work will focus on incorporating immune responses and the natural production of DIPs into the mathematical model presented here but will require further carefully curated data to assist in parameter estimation.

## Acknowledgments

G.B. and A.M. were supported by the Russian Science Foundation (research project number 23-11-00116). D.G. was financed by the Ministry of Science and Higher Education of the Russian Federation within the framework of state support for the creation and development of World-Class Research Centers “Digital biodesign and personalized healthcare” No. 075-15-2022-304. This manuscript has been internally reviewed at Los Alamos National Laboratory, and assigned the reference number LA-UR-22-29726 (CMP).

## Supporting information

**Figure S1.**
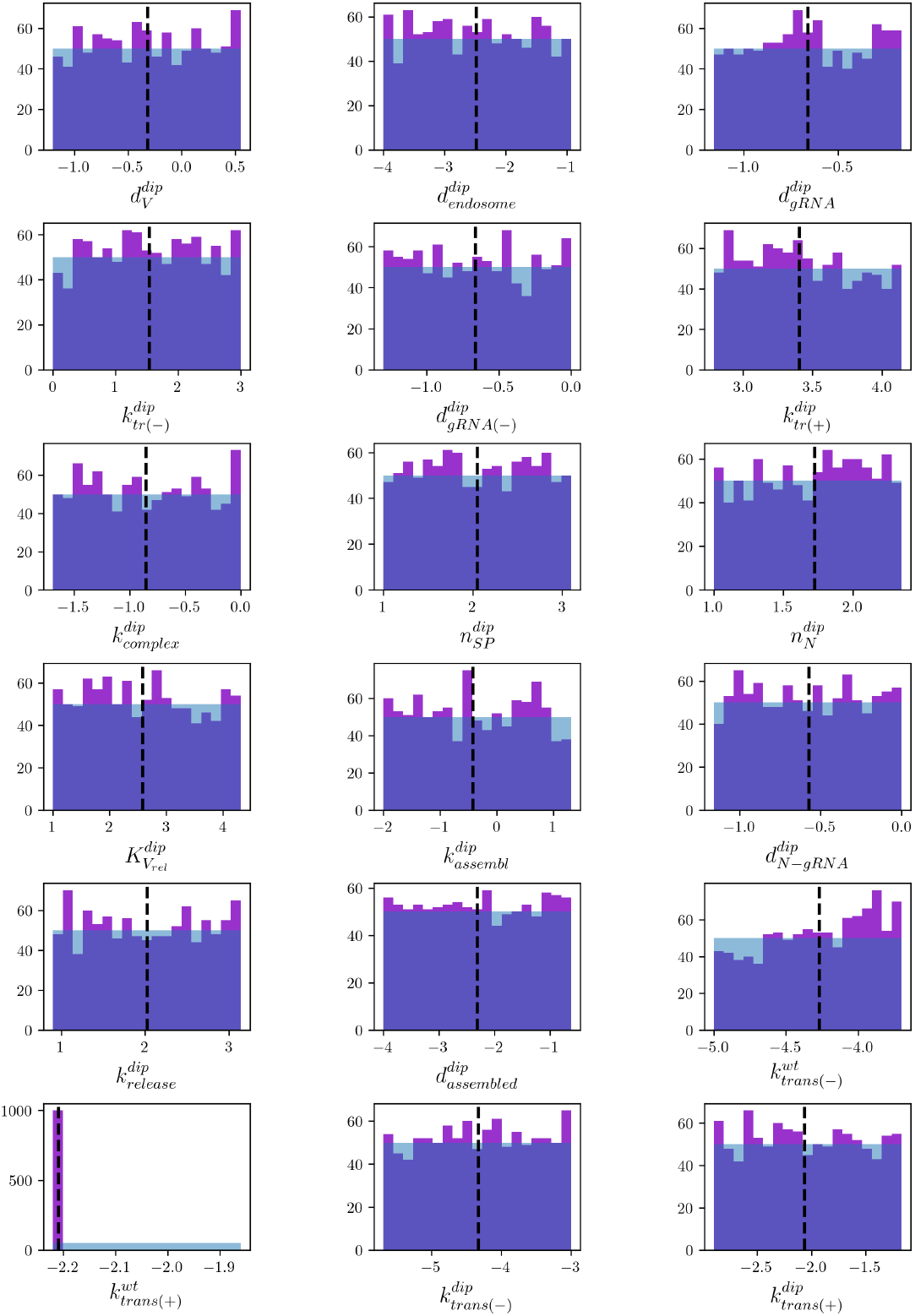
Posterior histograms. Posterior histograms of the top 0.1% sampled parameter sets from a total of 10^6^ accepted sets. Table 4 lists the search ranges used to obtain the above posterior histograms. **(Purple histogram:)** Posterior histogram of accepted parameter sets, **(blue histogram:)** histogram of prior beliefs, and **(black dashed line:)** the median parameter value listed in Table 4 used to generate Figures 4-9.

**Figure S2.**
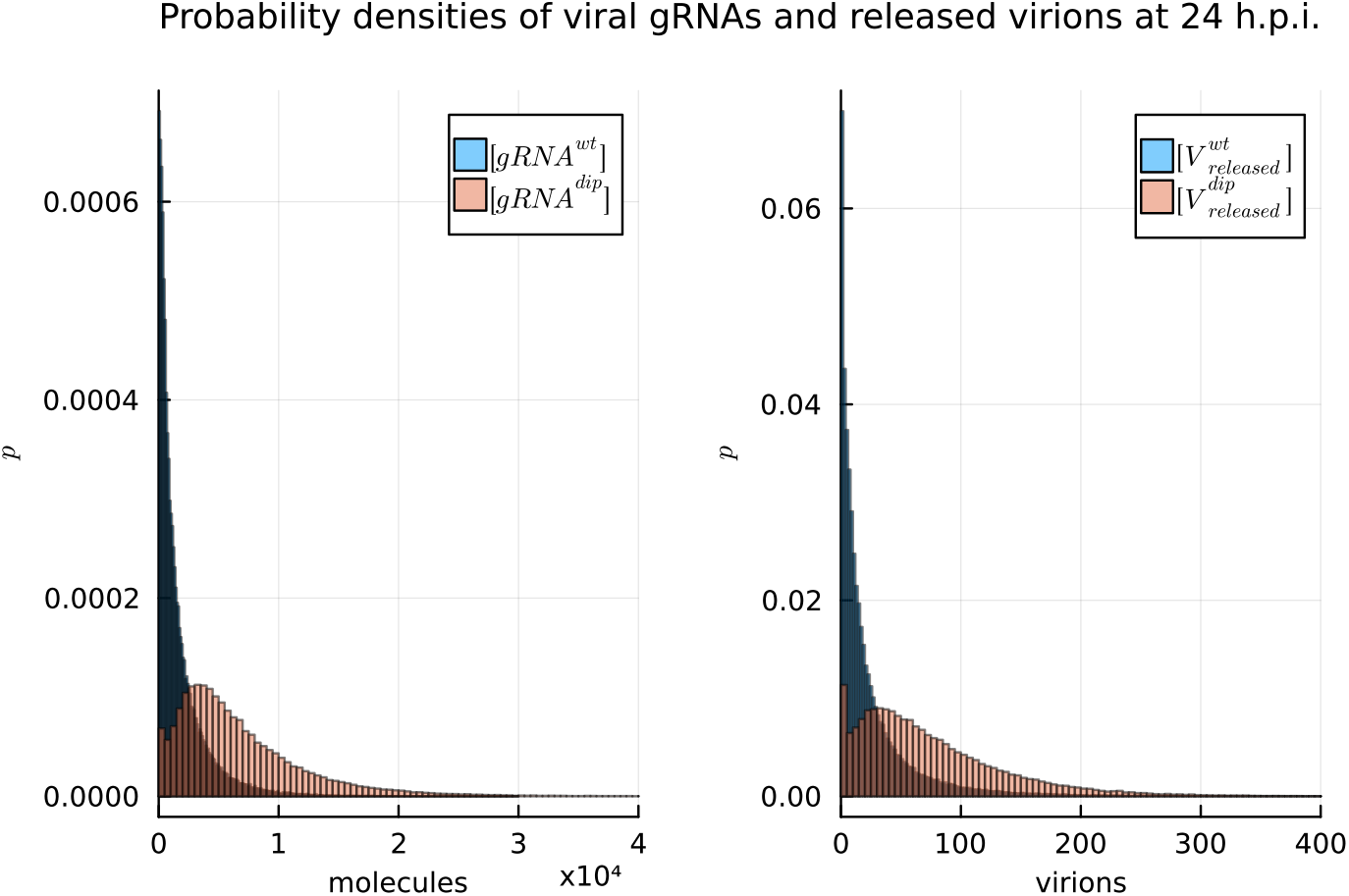
Stochastic model outputs 24 hours post-infection. The histograms for the numbers of **(left)** genomic RNA ([*gRNA*^*wt*^](24), [*gRNA*^*dip*^](24)) and **(right)** produced virions 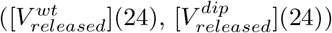 are shown for an ensemble of 10^6^ stochastic simulations with initial doses 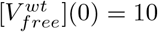 and 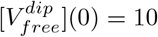. The histograms are normalised to approximate a true probability density distribution.

**Figure S3.**
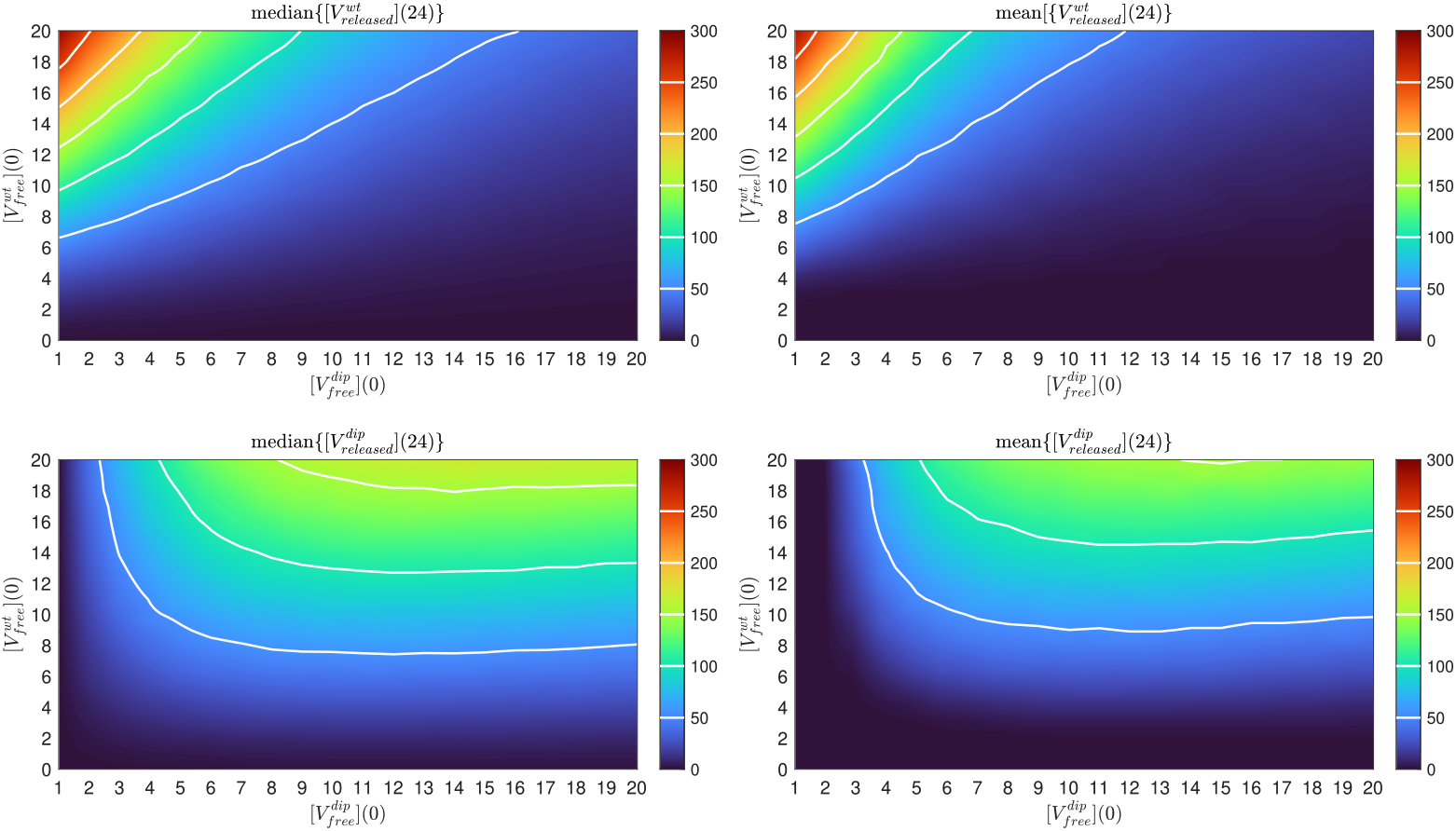
Effects of varying initial dose on viral particle release (as predicted by the stochastic model). **Left panels:** Median values, **right panels:** mean values are presented as the outputs of ensembles of 10^5^ trajectories simulated for each combination of the initial conditions. **Top:** Total WT virions released over the 24 hours post-infection for varying initial conditions of free WT virions 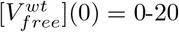 and free DIPs 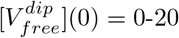 from the stochastic model. **Bottom:** Total DIP particles released for varying initial doses. The isolines shown on the heatmaps as white lines coincide with the corresponding ticks in the colorbars.

**Figure S4.**
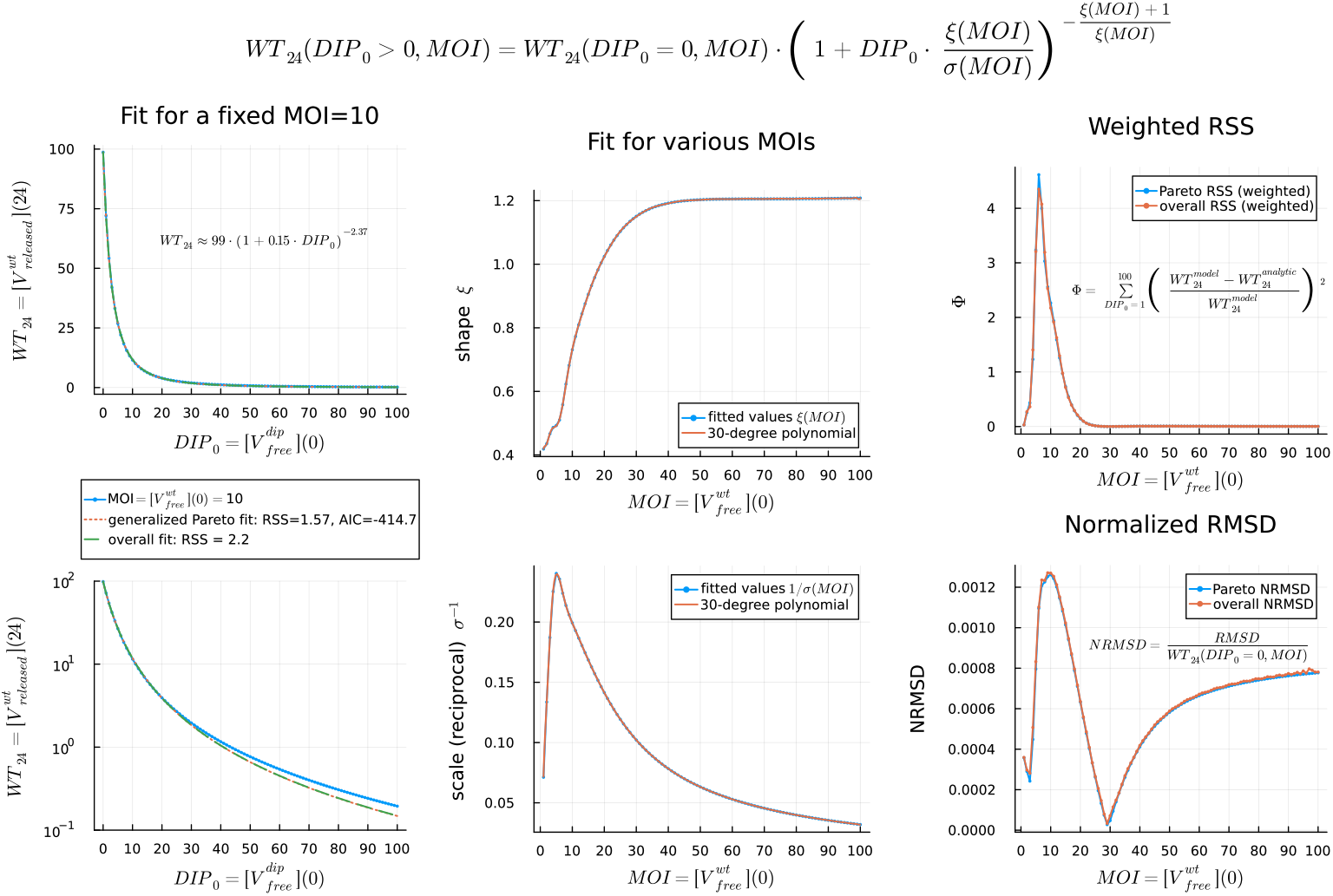
Fitting WT virion production as function of initial DIP doses with analytic expression. **Left:** fit of the deterministic model output (blue lines) for fixed *MOI* = 10 using Eq. (28), which is based on the probability density function of a generalised Pareto distribution (red dotted and green dashed lines). The upper plot is presented in linear scale, the lower one in logarithmic scale. The formula with estimated parameters is shown in the annotation of the upper plot. **Center:** fit of the parameters of the analytic expression (28) for various MOIs. The upper plot shows the fitted values of the parameter *ξ* and the lower one the reciprocal of the parameter *σ*. The fitted estimates *ξ*(*MOI*) and 1*/σ*(*MOI*) can be approximated closely with a 30-degree Chebyshev polynomial (red lines). The *overall fit* in the left panel denotes the fit with Eq. (28), where the polynomials are used as *ξ*(*MOI*) and 1*/σ*(*MOI*). **Right:** the error of the fit with Eq. (28) for various MOIs. The upper plot shows the residual sum of squares (RSS) weighted by the data values for each *DIP*_0_. The lower plot shows the root-mean-square deviation (RMSD) normalised by the number of produced WT virions with *DIP*_0_ = 0.

**Figure S5.**
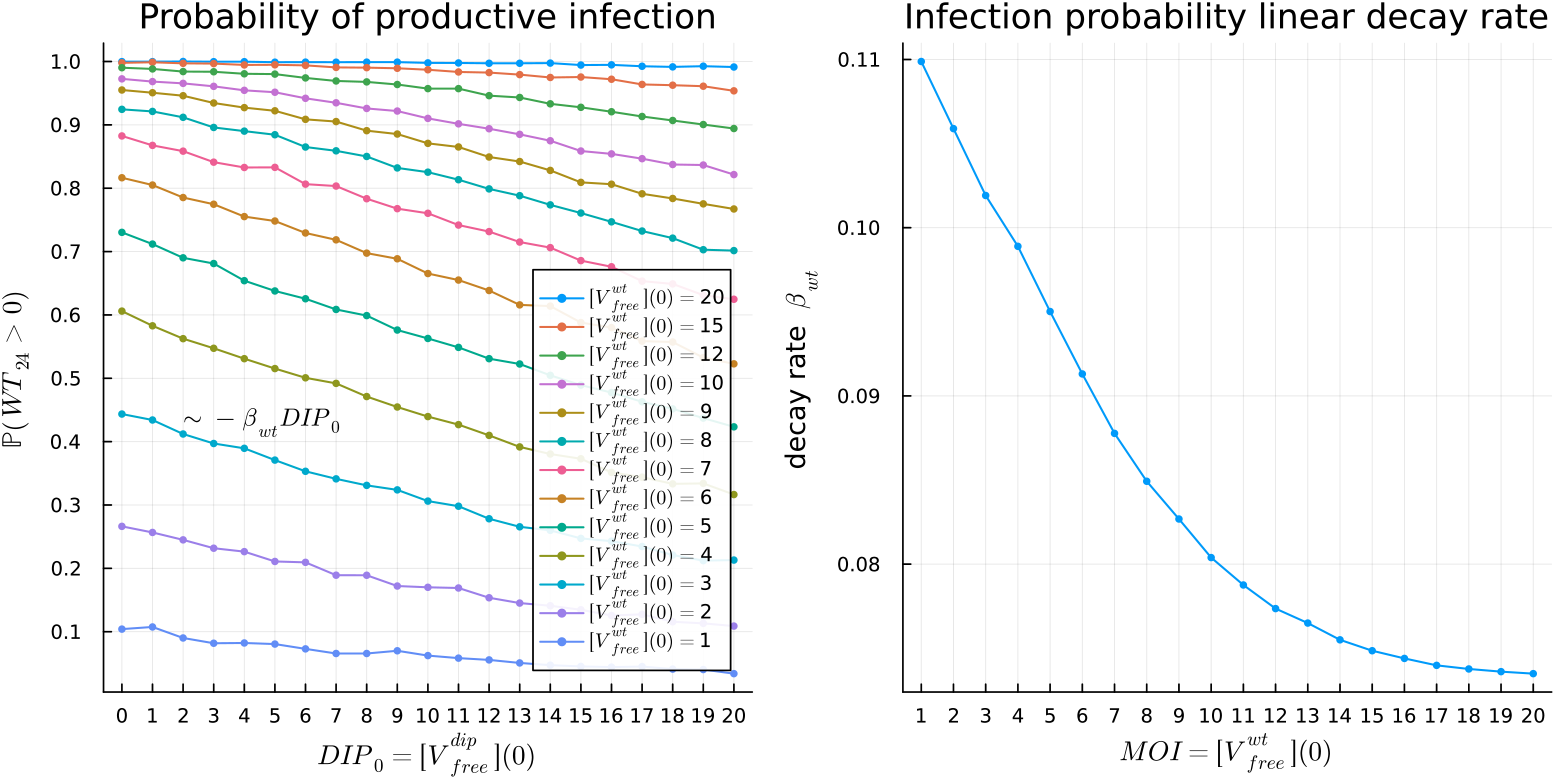
The estimation of the effect of DIP dose on the probability of productive infection as a function of MOI. **Left:** the effect of DIP initial dose on the probability of productive infection. For each MOI, the probability decreases linearly as DIP dose, *DIP*_0_, increases. **Right:** the dependence of the linear decay rate, *β*_*wt*_, on MOI. The decay rate, and therefore the effect of DIP dose, decrease with increase in MOI.

## References

1. Wu D, Wu T, Liu Q, Yang Z. The SARS-CoV-2 outbreak: what we know. International Journal of Infectious Diseases. 2020;94:44–48.

2. Li X, Wang W, Zhao X, Zai J, Zhao Q, Li Y, et al. Transmission dynamics and evolutionary history of 2019-nCoV. Journal of medical virology. 2020;92(5):501–511.

3. Martellucci CA, Flacco ME, Cappadona R, Bravi F, Mantovani L, Manzoli L. SARS-CoV-2 pandemic: An overview. Advances in biological regulation. 2020;77:100736.

4. Organization WH, et al. Coronavirus disease 2019 (COVID-19): situation report, 73; 2020.

5. Colson P, Rolain JM, Raoult D. Chloroquine for the 2019 novel coronavirus SARS-CoV-2; 2020.

6. Morse JS, Lalonde T, Xu S, Liu WR. Learning from the past: possible urgent prevention and treatment options for severe acute respiratory infections caused by 2019-nCoV. Chembiochem. 2020;21(5):730–738.

7. Castells MC, Phillips EJ. Maintaining safety with SARS-CoV-2 vaccines. New England Journal of Medicine. 2021;384(7):643–649.

8. Boehm E, Kronig I, Neher RA, Eckerle I, Vetter P, Kaiser L, et al. Novel SARS-CoV-2 variants: the pandemics within the pandemic. Clinical Microbiology and Infection. 2021;27(8):1109–1117.

9. Jangra S, Ye C, Rathnasinghe R, Stadlbauer D, Alshammary H, Amoako AA, et al. SARS-CoV-2 spike E484K mutation reduces antibody neutralisation. The Lancet Microbe. 2021;2(7):e283–e284.

10. Naqvi AAT, Fatima K, Mohammad T, Fatima U, Singh IK, Singh A, et al. Insights into SARS-CoV-2 genome, structure, evolution, pathogenesis and therapies: Structural genomics approach. Biochimica et Biophysica Acta (BBA)-Molecular Basis of Disease. 2020;1866(10):165878.

11. Fu Y, Cheng Y, Wu Y. Understanding SARS-CoV-2-mediated inflammatory responses: from mechanisms to potential therapeutic tools. Virologica Sinica. 2020;35(3):266–271.

12. Alnaji FG, Brooke CB. Influenza virus DI particles: Defective interfering or delightfully interesting? PLoS pathogens. 2020;16(5):e1008436.

13. Fatehi F, Bingham RJ, Dechant PP, Stockley PG, Twarock R. Therapeutic interfering particles exploiting viral replication and assembly mechanisms show promising performance: a modelling study. Scientific reports. 2021;11(1):1–10.

14. Bdeir N, Arora P, Gärtner S, Hoffmann M, Reichl U, Pöhlmann S, et al. A system for production of defective interfering particles in the absence of infectious influenza A virus. PLoS One. 2019;14(3):e0212757.

15. Chaturvedi S, Vasen G, Pablo M, Chen X, Beutler N, Kumar A, et al. Identification of a therapeutic interfering particle—A single-dose SARS-CoV-2 antiviral intervention with a high barrier to resistance. Cell. 2021;184(25):6022–6036.

16. Smither SJ, Garcia-Dorival I, Eastaugh L, Findlay JS, O’Brien LM, Carruthers J, et al. An Investigation of the Effect of Transfected Defective, Ebola Virus Genomes on Ebola Replication. Frontiers in cellular and infection microbiology. 2020;10:159.

17. Rouzine IM, Weinberger LS. Design requirements for interfering particles to maintain coadaptive stability with HIV-1. Journal of virology. 2013;87(4):2081–2093.

18. Grebennikov D, Karsonova A, Loguinova M, Casella V, Meyerhans A, Bocharov G. Predicting the Kinetic Coordination of Immune Response Dynamics in SARS-CoV-2 Infection: Implications for Disease Pathogenesis. Mathematics. 2022;10(17):3154. doi:10.3390/math10173154.

19. Perelson AS, Ke R. Mechanistic modeling of SARS-CoV-2 and other infectious diseases and the effects of therapeutics. Clinical Pharmacology & Therapeutics. 2021;109(4):829–840.

20. Zhao Y, Xing Y. A delayed dynamical model for COVID-19 therapy with defective interfering particles and artificial antibodies. Discrete & Continuous Dynamical Systems-B. 2021;.

21. Grebennikov D, Kholodareva E, Sazonov I, Karsonova A, Meyerhans A, Bocharov G. Intracellular Life Cycle Kinetics of SARS-CoV-2 Predicted Using Mathematical Modelling. Viruses. 2021;13(9). doi:10.3390/v13091735.

22. Zhang XY, Trame MN, Lesko L J, Schmidt S. Sobol Sensitivity Analysis: A Tool to Guide the Development and Evaluation of Systems Pharmacology Models. CPT: Pharmacometrics and Systems Pharmacology. 25th February 2015;(2):69–79.

23. Sobol IM. Sensitivity analysis for non-linear mathematical models. Mathematical modelling and computational experiment. 1993;1:407–414.

24. Ozono S, Zhang Y, Ode H, Sano K, Tan TS, Imai K, et al. SARS-CoV-2 D614G spike mutation increases entry efficiency with enhanced ACE2-binding affinity. Nature communications. 2021;12(1):1–9.

25. Walls AC, Park YJ, Tortorici MA, Wall A, McGuire AT, Veesler D. Structure, function, and antigenicity of the SARS-CoV-2 spike glycoprotein. Cell. 2020;181(2):281–292.

26. Baggen J, Persoons L, Vanstreels E, Jansen S, Van Looveren D, Boeckx B, et al. Genome-wide CRISPR screening identifies TMEM106B as a proviral host factor for SARS-CoV-2. Nature Genetics. 2021;53(4):435–444.

27. Bocharov G, Romanyukha A. Mathematical model of antiviral immune response III. Influenza A virus infection. Journal of Theoretical Biology. 1994;167(4):323–360.

28. Baccam P, Beauchemin C, Macken CA, Hayden FG, Perelson AS. Kinetics of influenza A virus infection in humans. Journal of virology. 2006;80(15):7590–7599.

29. Zhu Y, Yu D, Yan H, Chong H, He Y. Design of potent membrane fusion inhibitors against SARS-CoV-2, an emerging coronavirus with high fusogenic activity. Journal of virology. 2020;94(14):e00635–20.

30. Heldt FS, Kupke SY, Dorl S, Reichl U, Frensing T. Single-cell analysis and stochastic modelling unveil large cell-to-cell variability in influenza A virus infection. Nature communications. 2015;6(1):1–12.

31. Irigoyen N, Firth AE, Jones JD, Chung BYW, Siddell SG, Brierley I. High-resolution analysis of coronavirus gene expression by RNA sequencing and ribosome profiling. PLoS pathogens. 2016;12(2):e1005473.

32. Buccitelli C, Selbach M. mRNAs, proteins and the emerging principles of gene expression control. Nature Reviews Genetics. 2020;21(10):630–644.

33. Kim D, Lee JY, Yang JS, Kim JW, Kim VN, Chang H. The architecture of SARS-CoV-2 transcriptome. Cell. 2020;181(4):914–921.

34. Gasteiger E, Gattiker A, Hoogland C, Ivanyi I, Appel RD, Bairoch A. ExPASy: the proteomics server for in-depth protein knowledge and analysis. Nucleic acids research. 2003;31(13):3784–3788.

35. Nelson DL, Lehninger AL, Cox MM. Lehninger principles of biochemistry. Macmillan; 2008.

36. Adelman K, La Porta A, Santangelo TJ, Lis JT, Roberts JW, Wang MD. Single molecule analysis of RNA polymerase elongation reveals uniform kinetic behavior. Proceedings of the National Academy of Sciences. 2002;99(21):13538–13543.

37. Klein S, Cortese M, Winter SL, Wachsmuth-Melm M, Neufeldt CJ, Cerikan B, et al. SARS-CoV-2 structure and replication characterized by in situ cryo-electron tomography. Nature communications. 2020;11(1):1–10.

38. Zinzula L, Basquin J, Bohn S, Beck F, Klumpe S, Pfeifer G, et al. High-resolution structure and biophysical characterization of the nucleocapsid phosphoprotein dimerization domain from the Covid-19 severe acute respiratory syndrome coronavirus 2. Biochemical and biophysical research communications. 2021;538:54–62.

39. Chen IJ, Yuann JMP, Chang YM, Lin SY, Zhao J, Perlman S, et al. Crystal structure-based exploration of the important role of Arg106 in the RNA-binding domain of human coronavirus OC43 nucleocapsid protein. Biochimica et Biophysica Acta (BBA)-Proteins and Proteomics. 2013;1834(6):1054–1062.

40. Spencer KA, Hiscox JA. Characterisation of the RNA binding properties of the coronavirus infectious bronchitis virus nucleocapsid protein amino-terminal region. FEBS letters. 2006;580(25):5993–5998.

41. Spencer KA, Dee M, Britton P, Hiscox JA. Role of phosphorylation clusters in the biology of the coronavirus infectious bronchitis virus nucleocapsid protein. Virology. 2008;370(2):373–381.

42. Bar-On YM, Flamholz A, Phillips R, Milo R. Science Forum: SARS-CoV-2 (COVID-19) by the numbers. elife. 2020;9:e57309.

43. Jack A, Ferro LS, Trnka MJ, Wehri E, Nadgir A, Nguyenla X, et al. SARS-CoV-2 nucleocapsid protein forms condensates with viral genomic RNA. PLoS Biology. 2021;19(10):e3001425.

44. Cubuk J, Alston JJ, Incicco JJ, Singh S, Stuchell-Brereton MD, Ward MD, et al. The SARS-CoV-2 nucleocapsid protein is dynamic, disordered, and phase separates with RNA. Nature communications. 2021;12(1):1–17.

45. Viehweger A, Krautwurst S, Lamkiewicz K, Madhugiri R, Ziebuhr J, Hölzer M, et al. Direct RNA nanopore sequencing of full-length coronavirus genomes provides novel insights into structural variants and enables modification analysis. Genome research. 2019;29(9):1545–1554.

46. Neuman BW, Kiss G, Kunding AH, Bhella D, Baksh MF, Connelly S, et al. A structural analysis of M protein in coronavirus assembly and morphology. Journal of structural biology. 2011;174(1):11–22.

47. Yao H, Song Y, Chen Y, Wu N, Xu J, Sun C, et al. Molecular architecture of the SARS-CoV-2 virus. Cell. 2020;183(3):730–738.

48. Gordon DE, Hiatt J, Bouhaddou M, Rezelj VV, Ulferts S, Braberg H, et al. Comparative host-coronavirus protein interaction networks reveal pan-viral disease mechanisms. Science. 2020;370(6521).

49. Shcherbatova O, Grebennikov D, Sazonov I, Meyerhans A, Bocharov G. Modeling of the HIV-1 life cycle in productively infected cells to predict novel therapeutic targets. Pathogens. 2020;9(4):255.

50. Mooney J, Thakur S, Kahng P, Trapani JG, Poccia D. Quantification of exocytosis kinetics by DIC image analysis of cortical lawns. Journal of chemical biology. 2014;7(2):43–55.

51. Toni T, Welch D, Strelkowa N, Ipsen A, Stumpf MP. Approximate Bayesian computation scheme for parameter inference and model selection in dynamical systems. Journal of the Royal Society Interface. 2009;6(31):187–202.

52. Gillespie DT. Stochastic Simulation of Chemical Kinetics. Annual Review of Physical Chemistry. 2007;58(1):35–55. doi:10.1146/annurev.physchem.58.032806.104637.

53. Marchetti L, Priami C, Thanh VH. Simulation Algorithms for Computational Systems Biology. Texts in Theoretical Computer Science. An EATCS Series. Cham: Springer International Publishing; 2017. Available from: http://link.springer.com/10.1007/978-3-319-63113-4.

54. Sazonov I, Grebennikov D, Meyerhans A, Bocharov G. Markov Chain-Based Stochastic Modelling of HIV-1 Life Cycle in a CD4 T Cell. Mathematics. 2021;9(17):2025. doi:10.3390/math9172025.

55. Chaturvedi S, Vasen G, Pablo M, Chen X, Beutler N, Kumar A, et al. Identification of a therapeutic interfering particle—A single-dose SARS-CoV-2 antiviral intervention with a high barrier to resistance. Cell. 2021;184(25):6022–6036.e18. doi:10.1016/j.cell.2021.11.004.

56. V’kovski P, Kratzel A, Steiner S, Stalder H, Thiel V. Coronavirus biology and replication: implications for SARS-CoV-2. Nature Reviews Microbiology. 2021;19(3):155–170. doi:10.1038/s41579-020-00468-6.

57. Mendonça L, Howe A, Gilchrist JB, Sheng Y, Sun D, Knight ML, et al. Correlative multi-scale cryo-imaging unveils SARS-CoV-2 assembly and egress. Nature Communications. 2021;12(1):1–10.

58. Sazonov I, Grebennikov D, Meyerhans A, Bocharov G. Sensitivity of SARS-CoV-2 Life Cycle to IFN Effects and ACE2 Binding Unveiled with a Stochastic Model. Viruses. 2022;14(2):403. doi:10.3390/v14020403.

59. Rand U, Kupke SY, Shkarlet H, Hein MD, Hirsch T, Marichal-Gallardo P, et al. Antiviral activity of influenza A virus defective interfering particles against SARS-CoV-2 replication in vitro through stimulation of innate immunity. bioRxiv. 2021;.

60. Roux L, Simon AE, Holland JJ. Effects of defective interfering viruses on virus replication and pathogenesis in vitro and in vivo. Advances in virus research. 1991;40:181–211.

61. Li D, Lin MH, Rawle DJ, Jin H, Wu Z, Wang L, et al. Dengue virus-free defective interfering particles have potent and broad anti-dengue virus activity. Communications biology. 2021;4(1):1–11.

62. Dempsey P, Vaidya S, Cheng G. The art of war: Innate and adaptive immune responses. Cellular and Molecular Life Sciences CMLS. 2003;60(12):2604–2621.

63. tenOever B. The Evolution of Antiviral Defense Systems. Cell Host & Microbe. 2016;19(2):142–149. doi:10.1016/j.chom.2016.01.006.

64. Katze MG, He Y, Gale M. Viruses and interferon: a fight for supremacy. Nature Reviews Immunology. 2002;2(9):675–687.

